# Revisiting the role of *Acinetobacter* spp. in EBPR systems

**DOI:** 10.1101/2023.04.01.535225

**Authors:** Yuan Yan, IL Hana, Jangho Lee, Guangyu Li, Varun Srinivasan, Kester McCullough, Stephanie Klaus, Da Kang, DongQi Wang, Anand Patel, Jim McQuarrie, Beverley M. Stinson, Christine deBarbadillo, Paul Dombrowski, Charles Bott, April Z. Gu

**Author notes:** Joint first authors (Yuan Yan, IL Hana). Corresponding authors: April Gu.

## Abstract

Side-stream Enhanced biological phosphorus removal (S2EBPR) has been incorporated with B-stage process to enable simultaneous phosphorus and nitrogen removal. However, the dominating phosphorus accumulating organisms (PAOs) in this novel configuration has not been evaluated. The dominance of Acinetobacter was confirmed by 16S sequencing. In addition, single cell Raman spectrum (SCRS) analysis in couple with in situ fluorescence in situ hybridization (FISH) was applied to obtain the feature spectrum and verify the phosphorus release/uptake activity of Acinetobacter spp. The phenotypic profiling further suggested the dominance of Acinetobacter-like organisms among all poly-phosphorus containing organisms and only certain phenotypic Acinetobacter (oligotype 1) contribution to P-removal in a unique HRAS-P(D)N-S2EBPR system. The findings suggest that Acinetobacter may outcompete other heterotrophic organisms in EBPR systems due to their sensitivity to operational conditions. However, stable P-removal was only observed during a specific section of the operation period, coinciding with an increase in the VFA/P ratio. Further research is needed to identify the phenotypes of Acinetobacter responsible for P-removal in EBPR systems. The study contributes to a better understanding of the microbial ecology and engineering aspects of EBPR systems and wastewater treatment in general.

## Highlights

- Sufficient (85%) phosphorus removal was achieved (Eff. 0.63 ± 0.58 mg-P L^−1^) with low abundance of known PAOs (0.6 ± 0.3%)
- *Acinetobacter* dominated (median 16%) the microbial community and was proven to have PAO activity by FISH-Raman techniques
- Combined 16S rRNA-gene based amplicon sequencing and oligotyping analysis revealed subgroup of Acinetobacter (oligo1) might contribute to the phosphorus removal

## 1. Introduction

Biological nutrient removal (BNR) via nitrification and denitrification is widely used to control wastewater nitrogen. However, the conventional methods have concerns in energy consumption, sludge production, and greenhouse gas emission (Li et al., 2018; Zhang et al., 2020). The Adsorption-Biooxidation (A-B) processes, where A-stage diverts most organic matter to a digester for energy recovery, and B-stage aims for carbon and energy-efficient nutrient removal (Miller et al., 2017), is among several approaches proposed towards more self-sufficient wastewater treatment. In the A-B processes, partial nitrification (or denitrification)/anammox (P(D)N/A) that can achieve N-removal with less resource and sludge production is often used for the B-stage (Zhang et al., 2020). However, the incorporation of enhanced biological phosphorus removal (EBPR) into the A-B stage process has not been targeted previously because presumed inherent incompatibility with N-removal processes. The favorable conditions for the phosphorus accumulating organisms (PAOs), such as adequate C/P ratio (rbCOD/P > 25), carbon content, and cycling anaerobic-aerobic (An/Aer) condition, are crucial for stable P-removal in EBPR but conflict with those preferred in P(D)N/A process. For instance, P(D)N/A process requires restricted DO or intermittent aeration to suppress NOB (Lackner and Horn, 2013), low COD to prevent heterotroph inhibition on autotrophic nitrification community (Wang et al., 2019). This conditions are inconsistent with those preferred for optimization of EBPR, particularly in handling the weak C/P ratio exaggerated in the B-stage processes (Gu et al., 2008; Neethling et al., 2006).

The recently developed side-stream EBPR (S2EBPR) process, which involves ferment/hydrolysis of a portion of return activated sludge (RAS, 10-40%) in a side-stream reactor to foster PAO enrichment, helps overcome the reliance on influent condition (Gu et al., 2018; Onnis-Hayden et al., 2020; D. Wang et al., 2019). This approach could be a viable option for the P(D)N/A process to incorporate bio-P removal as it creates complex VFA composition and kinetic conditions favorable for the enrichment of PAOs. The direct integration of EPBR to P(D)N/A process is expected to affect both N-removal and P-removal performances. The microbial populations in the unique A-B process coupled with S2EBPR are subject to selection forces that are different from conventional EBPR or A-B processes without S2EBPR, especially the identities of PAO populations remain largely unknown.

Various candidate PAOs, including Candidatus *Accumulibacter*, *Tetrasphaera* and *Dechloromonas* are consistently reported to be important for the performance of full-scale EBPR processes (Stokholm-Bjerregaard et al., 2017; Fernando et al., 2019). Other than that, *Acinetobacter* was originally perceived as the primary PAO responsible for P-removal because they are the dominant poly-P accumulating isolates found in many EBPR facilities (Fuhs and Chen, 1975). Later, Wagner et al., (1994) found poly-P inclusion in subclass of Proteobacteria in sludge had a higher relative abundance than *Acinetobacter* through *in situ* fluorescent staining, indicating that culture-dependent isolation method favors *Acinetobacter* growth and causes an overestimation of their role in EBPR processes. Furthermore, Saunders et al. (2016) excluded *Acinetobacter* from the frequently abundant (core) OTUs that are important for WWTPs across Denmark, as their abundance was owing to the high immigration from the influent rather than the proliferation in treatment processes. Despite this, a recent study analyzed the global diversity of bacterial community in WWTPs (269 WWTPs in 23 countries on six continents) and defined *Acinetobacter* as a core OTU with positive Spearman’s rank correlation with TP removal rate (Wu et al., 2019). In addition, P-removal by *Acinetobacter* are reported in other WWTPs (Han et al., 2018; Liu et al., 2014), making their contribution to EBPR performance ambiguous.

Recently, our team reported a pilot-scale integration of S2EBPR into a A-B stage process (HRAS-P(D)N-S2EBPR) located at the Hampton Road Sanitation District (HRSD) Chesapeake-Elizabeth Plant in Virginia Beach, VA (Kang et al., 2023; Klaus et al., 2019; McCullough et al., 2020). In this study, the performance and the microbial communities of this pilot plant were monitored and analyzed for one year. A period of low effluent-P (PO_4_^−^-P < 1.0 mg-P L^−1^) was achieved with microbial community dominated by *Acinetobacter* (median 16%) and very low conventional PAO abundance (0.05 – 1.2%) ( Klaus, 2019). This observation motivated us to further investigate the roles and involvements of *Acinetobacter* in HRAS-P(D)N-S2EBPR. The application of Raman micro-spectroscopy has opened new opportunities to investigate phenotypical characteristics of individual microorganisms. This technique has been used to separate and classify the microbial groups important for the EBPR process by hierarchical clustering analysis of Raman spectral fingerprints (Li et al., 2018); and to track changes in intracellular polymers, such as poly-P, PHA and glycogen (Majed and Gu, 2010, p. 20) in specific microbial genus by combining with cytogenetic technique, such as fluorescence in situ hybridization (FISH) (Fernando et al., 2019). Throughout the operation period, phylogenetic and phenotypic characteristics of microorganisms were analyzed with 16S rRNA amplicon-based analysis as well as the Raman/FISH-Raman micro-spectroscopy. The results provide evidence that *Acinetobacter* dominated and contributed to phosphorus removal in the HRAS-P(D)N/A-S2EPBR system.

## 2. Material and Methods

### 2.1. Pilot Plant Operation Condition

Configuration and performance of the HRAS-P(D)N-S2EBPR pilot plant are described by (Klaus, 2019). Briefly, the pilot plant was in A-B processes configuration (Figure S1). The A-stage was an HRAS process receiving screened (2 mm) raw wastewater at 20 °C (HRT 45 min, SRT 8 h). The B-stage was an intermittently aerated tank receiving A-stage clarifier effluent for P(D)N (HRT 4 – 5h; SRT ca. 8 d without S2EBPR tank; aerobic SRT 3 – 5 d). The S2EBPR tank received a fraction of RAS (ca. 10 – 30%) diverted from the secondary clarifier for fermentation and PAOs enrichment. It also received fermentate from the A-stage fermenter for performance boost. Effluents from the A-stage clarifier and S2EBPR tank was briefly mixed in a tank before entering the B-stage. The performance of the studied system was monitored by measuring MLSS, MLVSS, sCOD, PO_4_^−^-P, NH_4_ ^+^-N, NO_2_^−^-N, and NO_3_^−^-N (American Public Health Association and others, 2005).

### 2.2. EBPR activity test

Periodic routine and detailed P-release/uptake tests were performed on the studied system to measure PAO and EBPR activity. These tests were conducted following the procedure described by Gu et al., (2008). Briefly, periodic routine tests were performed two to four times a month. In these tests, fresh activated sludge from the CSTR4 was transferred to a batch reactor and supplemented with sodium acetate (HAc) to a concentration of 80 mg-HAc L^−1^. Anaerobic condition (DO < 0.01 mg L^−1^) was created for 60 min by purging the sludge with N_2_ gas and PAO activity related P-release and carbon uptake was measured. Subsequently, aerobic condition (DO > 2.0 mg L^−1^) was created for 60 min by purging the sludge with air and PAO activity related P-uptake was measured. Samples were collected every 30 min. for sCOD, PO_4_^−^ −P, NH_4_^+^-N, NO_2_^−^-N, and NO_3_^−^-N, measurements.

The detailed EBPR activity test was performed when effluent PO_4_^−^-P was < 1.0 mg-P L^−1^. In this test, HAc (130 mg L^−1^ as sCOD) and A-stage fermentate (180 mg L^−1^ as sCOD) were used as C-substrate, respectively. Supernatant samples were filtered with 0.45 um membrane every 30 min for PO_4_^−^-P, NH_4_ ^+^-N, NO_2_^−^-N, and NO_3_^−^-N measurements. Furthermore, sludge samples were collected for polyhydroxyalkanoate (PHA), glycogen (Gly), regular Raman and FISH-Raman analysis. PHA and Gly were analyzed following the methods described in previous studies (X. Wang et al., 2019). Briefly, PHA was extracted from lyophilized sludge using acidified methanol (3% sulfuric acid) containing chloroform and analyzed using a gas chromatography-mass spectrometry (Agilent, USA). Glycogen in sludge was extracted in a similar way but using 0.9M HCl and analyzed using a liquid chromatography-mass spectrometer (Thermo Scientific, USA).

### 2.3. Sample preparation for Raman and FISH-Raman spectroscopy

Raman and FISH-Raman samples were prepared following the methods described in previous studies (Fernando et al., 2019; Li et al., 2018). Briefly, samples were collected at the start of anaerobic (t = 0 min), end of anaerobic (t = 60 min), and end of aerobic (t = 120 min) phase of a P-release/uptake test and washed three times with 0.9% NaCl. For the regular Raman, washed samples were diluted 100 times with 1 X PBS then disrupted using a 26-gauge needle and 1 mL syringe for uniform distribution of cells. Four separate 2.5 µL drops were dried on a CaF_2_ slide per sample (Crystran Ltd., Dorset, UK). For the FISH-Raman, samples from each time point were centrifuged at 6,000 rpm, and supernatants were discarded. Pellets were resuspended in ice-cold 4% PFA solution then left to fix in ice for 2 h shielded from light. Fixed samples were washed, diluted, and dried on a CaF_2_ slide as described above.

The ACA23a probe (5’-ATCCTCTCCCATACTCTA-3’) was used to target genes *Acinetobacter* (Wagner et al., 1994) for the FISH-Raman analysis. This probe had 18,224 hits on the RDP11 ProbeMatch tool (Cole et al., 2014), of which 98.29% are *Acinetobacter*. The probe covered 82.11% of the targeted genus, including those reported to accumulate poly-P, such as *A. tandoii* (Zhang et al., 2019), *A. junii* (Ren et al., 2014), and *A. calcoaceticus* (Hrenović et al., 2003). The hybridization was done following the protocol described previously (Fernando et al., 2019).

The Raman spectra was acquired using a Horiba Labram Evolution (Horiba, Kyoto, Japan) equipped with 50X objective (Olympus LMPLFLN 50x, Tokyo, Japan) following previously described protocol (Li et al., 2018). Excitation was provided by a 532 nm diode laser. The spectra were collected from 400 to 2,000 cm^−1^ with ND 25%, acquisition time 20 s, and accumulation 3.

### 2.4. Single cell Raman spectrum (SCRS) analysis and phenotypic profiling

The presence of polyphosphate (poly-P) in single cells was identified by the two signature peaks in the range of 685-715 cm^-1^ and 1150-1180 cm^-1^ based on previous study (Majed and Gu, 2010). And the second peak was used for quantification. The relative intensity of poly-P for single cell was normalized by the intensity of amide I vibration at 1640-1690 cm^-1^. For effective analysis, sufficient (78-137) bacterial Raman spectrum were collected according to Li et al., (2022).

Since the cell fixation and FISH staining process would lead to the loss of poly-P signals, we utilized combined Raman-spectral fingerprint and multivariate methods, Raman-phenotypic profiling, to separate cells with phenotypic feature similar to *Acinetobacter spp*. The Raman fingerprints of ACA23a hybridized cells were used as the SCRS reference. The detailed procedure were depicted in (Li et al., 2018). Briefly, the hierarchical clustering analysis (HCA) was applied on all the SCRSs from sludge samples and *Acinetobacter spp*. identified by FISH probe to obtain the phenotypic profiles. Spectra were first binned with a windows size of 5 to reduce the impact of peak shift, followed by HCA using square root cosine similarity (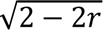, r-correlation efficient) as spectra pair-wise metric, average linkage, and 1.1 as cutoff threshold.

### 2.5. Microbial community analysis

#### 2.5.1. 16S rRNA amplicon-based identification of candidate PAOs

The microbial community of the HRAS-P(D)N-S2EBPR system was analyzed with 16S rRNA gene amplicon sequencing. Genomic DNA was extracted from sludge samples with using a Qiagen Power soil DNA extraction kit (Qiagen, Germany). University of Connecticut-MARS facility performed bacterial amplicon sequencing targeting V4 regions (primer: 515F 5’-GTGCCAGCMGCCGCGGTAA-3’, and 806R 5’-GGACTACHVGGGTWTCTAAT-3’) with Illumina MiSeq system. Acquired sequences were analyzed using Mothur (Schloss et al., 2009), and the paired-end reads were merged and clustered at a 97% similarity for genus-level classification. Representative OTUs for sequences were identified based on the Silva database (Quast et al., 2012). Non-metric multidimensional scaling (NMDS) with distance bray was plotted for the microbial community using the meta MDS function from the vegan package (R Core Team, 2013).

Five additional putative PAOs relevant to wastewater treatment systems were references based on MiDAs database (https://midasfieldguide.org/guide/search), namely Ca. *Accumulibacter* (Martín et al., 2006), *Tetrasphaera* (Kristiansen et al., 2013), *Arthrobacter* (Li et al., 2013), *Microlunatus* (Kawakoshi et al., 2012), *Obscuribacter* (Soo et al., 2014). Polyphosphate kinase I (*ppk*1), polyphosphate kinase II (*ppk*2) and exophosphatase (*ppx*) are considered key functional genes in poly-P metabolism. At the genus level, taxa were filtered against a list of reference bacterial genera containing at least one species with *ppk*/*ppx* annotation to find all potential PAOs involved in the P-removal process. The lists generated from the NCBI Gene database contained 920 and 883 unique genera potentially carrying *ppk* and *ppx* gene, respectively (accessed July 30, 2020). The genera from the OTU classification were extracted if shared by both *ppk* and *ppx* reference genera lists.

#### 2.5.2. Oligotyping of *Acinetobacter*

The identified *Acinetobacter* OTUs were further resolved into oligotypes for assessment of their temporal profiles by oligotyping (Varun paper on oligo in WR, Eren et al., 2013). This approach first locates the high-variation base positions in multiple sequence alignment of 16S contigs and resolves subtypes in accordance with these bases. Extra positions were thereafter manually added to refine the preliminary oligotyping results until all oligotypes contained no base positions with entropy greater than 0.2. This criterion is equivalent to that each position has a most dominant base type (A, T, G, C or gap) ≥ 99.0%. The representative sequences from the three most abundant oligotypes (contains ≥ 10% total read counts) were extracted and taxonomically identified by searching through NCBI’s rna_refseq database using BLASTN. Representative sequences were taking the majority of at least 67% of all aligned sequences (with redundancy) in each oligotype.

#### 2.5.3. potential Metabolic pathway mapping of *Acinetobacter*

The genomic capabilities of closely related genus *Acinetobacter* identified by 16S rRNA amplicon sequencing were assessed. Their reference assemblies were downloaded from NCBI database (accessed 2019-04-16). The genes were predicted by prodigal v2.6.3 (Hyatt et al., 2010) then annotated via BlastKOALA (Kanehisa et al., 2016). The KEGG MODULE database (Kanehisa, 2006) was further consulted for essential gene identification and pathway reconstruction. A pathway-level metabolic capability was identified positive if at least 60% of genes in the respective KEGG module are found.

## 3. Results

### 3.1. System performance of the HRAS-P(D)N-SBPR process

The raw influent contains around 529 ± 160 mg/L COD and 6.11 ± 0.85 mg-P/L total phosphorus (TP) with the BOD/TP ratio ranges between 28-40 mg/mg. The COD/BOD ratio in raw influent is 2.73 ± 1.01 mg/mg. The HRAS process captured 55% of influent COD as WAS, leaving lower soluble biodegradable COD entering B-stage process (22 ± 9.3 mg-sCOD/mg-P). The average BOD, sCOD, TP and PO_4_^−^-P concentrations in B-stage influent are summarized in Table S1. The effluent PO_4_^−^-P of S2EBPR facilities in North America ranged between 0.2 – 0.9 mg-P L^−1^ with 79 – 100% achieving low effluent-P (Onnis-Hayden et al., 2020). Though the HRAS-P(D)N-S2EBPR system in this study had slightly higher effluent-P, it achieved effluent PO_4_^−^-P < 1.0 mg-P L^−1^ (P-limit, hereafter) (Figure 1A, Table S1). The influent BOD/P ratio maintained around 18.6 ± 3.5 to 22 ± 9.3 g/g (Table S1, Figure 1C), which is lower than previous reported S2EBPR system (38.4:1-102:1), but close to the suggested minimal C/P ratio (15:1-25:1 g rbCOD/g P) for EBPR (Neethling et al., 2006; Onnis-Hayden et al., 2020; D. Wang et al., 2019). However, the BOD/P ratio did not show clear relationship with effluent phosphorus. Only between days 184 – 267, the effluent-P was 0.9 ± 0.6 mg-P L^−1^, and 68% met P-limit. The effluent-P showed a decreasing trend (ca. 0.028 mg-P day^−1^) after RAS split ramping up to 21% and met P-limit from days 184. The effluent P and P removal efficiency averaged to be 0.63 ± 0.58 mg-P L^−1^ and 85%, respectively, during the best performance (day 184 – 267) (Figure S2). The VFA mass flow from A-stage fermentate to S2EBPR was increased during this period from 68.8 ± 16.8 to 116.5 ± 17.6 g-COD/d, leading to an increasing VFA/P (g-COD/g-P) ratio from 4.9 ± 1.5 to 7.5 ± 2.5 (Figure 1B), and an increasing VFA/sCOD (g-COD/g-COD) from 31 ± 5% to 42 ± 10% (Figure 1D). However, P removal deteriorated after a pre-denitrification tank was employed and VFA mass flow decreased on day 268 (Figure 1A, 1B, and S2).

**Figure 1.**
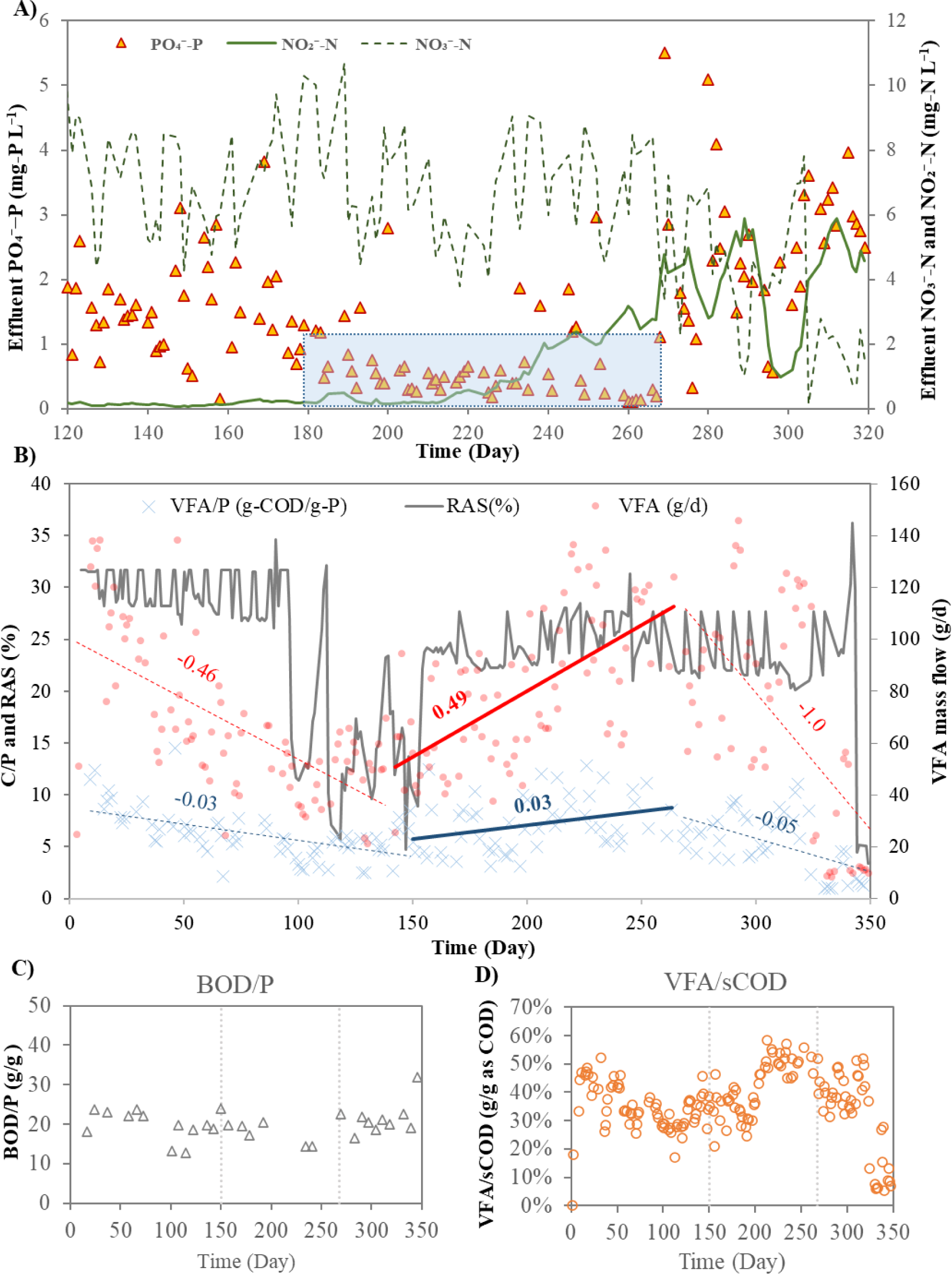
HRAS-P(D)N-S2EBPR performance. A) Nutrient content of the effluent. Measurements from day 0 – 120, and day 320 onwards are not shown. Shaded area represents period of PO4--P ˂ 1.0 mg-P L−1; B) VFA/P ratio, RAS split (%) and VFA mass flow (g-COD/d) into S2EBPR normalized to B-stage influent. Red and black solid-lines represents VFA and C/P trends at each period, respectively, C) BOD/P ratio in B-stage influent and D) the fermentate VFA normalized to the sCOD in B-stage influent.

The effluent NO_2_^−^-N was maintained below 0.63 mg-N L^−1^ for most the days during this study period until day 215 (Figure 1A). Stable nitrite accumulation occurred during day 216 to day 290 with highest concentration of 5.90 mg-N L^-1^. The periodic *ex-situ* nitrification rate test shows a decrease in the NOB/AOB activity ratio from 1.10 ± 0.11 to 0.91 ± 0.25, which did support the nitrite accumulation via partial nitrification pathway. The nitrite accumulation was disturbed on day 290, possibly from a brief operational issue with A-stage fermenter. The nitrite accumulation rebounded soon after and then lost again from day 313. During day 290-313, the NOB/AOB activity ratio decreased further to 0.75 ± 0.22. Also, onsite measurement of the P(D)N process by Klaus, (2019) suggested that the nitrite accumulation could be due to the partial-denitrification, indicating the mutual contribution of partial (de)nitrification on the nitrite accumulation.

It is interesting to see both nitrite accumulation and good phosphorus removal performance occurred at the same period (184-268) when the side-flow fermentate addition was high (Figure 1A).

### 3.2. EBPR metabolic activity and kinetics assessment

The PAO activity in the HRAS-P(D)N-S2EBPR system was monitored periodically with *ex-situ* P-release and P-uptake batch tests. When P-removal performance was good, the P-uptake and P-release rates were in the range of other facilities, corroborating moderate PAO activity (Table S2). The HAc-uptake rates were relatively consistent (*p* > 0.05) throughout the operation, implying relative stable activities of PAOs, glycogen accumulating organisms (GAOs), and OHOs. The P-release and HAc-uptake ratio (P-up/HAc) was 0.17 ± 0.03 P-mol/C-mol in the period of low effluent-P, which is lower than those from other facilities (0.38 – 0.54 mol-P/mol-C) as well as PAO enriched culture (0.55 mol-P/mol-C; average pH 7.4; Filipe et al., (2001)). The lower ratio suggests a relatively higher abundance of GAOs in the system. The competition for the C-substrate between PAOs and GAOs can impact P removal performance (Onnis-Hayden et al., 2020). Some level of PAO activity was also observed in during unstable performance. Interestingly, the PAO activity for the first 149 days were not statistically different (P > 0.05) from those after the P-removal destabilized, even though it had significantly lower effluent-P (*p* < 0.01). These results indicate the intrinsic capability to reach P-limit governed by the changes in PAO activities from the pilot operation.

In the PAO activity test, aerobic P-uptake/PHA-use (P/PHA) was 0.26 mol-P/mol-C using HAc as the C-source, which is lower than other facilities (0.7 ± 0.5 mol-P/mol-C) (Table S2). The ratio increased to 0.43 mol-P/mol-C when C-source is switch from acetate to the fermentate (combined HAc and propionate (HPr) fraction ca. 70% VFA; HAc/HPr ca. 1.1 mol-C/mol-C), suggesting the presence of PAOs that prefer variety of VFAs other than HAc. The anaerobic glycogen-use/HAc-uptake (Gly/HAc) was 0.50 mol-C/mol-C using HAc, which is higher than those of other facilities (0.15–0.35 mol-C/mol-C), consistent with the higher relative abundance of GAOs as discussed above. The ratio was 25% less using fermentate (0.34 mol-C/mol-C), being like other facilities. It was shown that S2EBPR enriched more diverse PAOs due to the more complex VFA composition in the S2EBPR in our study, in contrast to HAc dominant WW influent as in most cases. Higher level of HPr in relative to HAC was reported to favor PAOs over GAOs due to the lower specific HPr-uptake rate of GAOs than PAOs (Lopez-Vazquez et al., 2009; Oehmen et al., 2005). The overall efficient P removal activity achieved during 150-267 days in this study period demonstrates the success incorporation of S2EBPR into the HRAS-P(D)N-S2EBPR that enables simultaneous P removal and short-cut N removal.

### 3.3. Microbial community analysis: *Acinetobacter* is the major potential PAO

#### 3.3.1. Significantly low abundance of known PAOs over the studied period

The average richness (R) and Shannan index (H’) of the microbial community did not show significant change over the operation period (Table S3). Similarly, the NMDS analysis could not clearly separate the microbial community from different time points (Figure S3). These results suggest a relatively stable microbial community structure regardless of the changing P-removal performance. Based on MiDas database, PAOs (known PAOs including *Tetrasphaera*, Ca. Accumulibacter, and *Microlunatus*; putative PAO *Dechloromonas*) abundances were only 0.6 ± 0.3% during the investigation period. There were only two minor shifts over the studied period. The first was after the S2EBPR setup (before day 0), and the second was between day 300 – 320 (possibly with the removal of the pre-denitrification tank). After the incorporation of S2EBPR tank, the total known PAOs and GAOs dropped from 4.0% and 0.6% to 0.3% and 0.1%, respectively (Figure S4). Although total known PAO reached 2.1% at one point (day 231), the relative abundances of Ca. *Accumulibacter* and *Tetrasphaera* (0.04 ± 0.10% and 0.47 ± 0.32%) remained low throughout the operation period. Onnis-Hayden et al., (2020) reported a higher Ca. *Accumulibacter* (0.54 ± 0.06%) and *Tetrasphaera* (1.81 ± 0.1%) abundance in S2EBPR facilities compared to the studied system. However, this was still significantly less than the average PAO abundance (13.9%) found in the conventional EBPR facilities (Stokholm-Bjerregaard et al., 2017). Though low PAO abundance could partly be due to the poor Ca. *Accumulibacter* coverage of 16S rRNA amplicon analysis (Onnis-Hayden et al., 2020; D. Wang et al., 2019), known PAOs did not seem to be the dominant PAO in S2EBPR system.

#### 3.3.2. Genus *Acinetobacter*: Potential PAOs in the PN-S2EPBR system

We further evaluated the genera that possess ppk/ppx genes and found 78 genera in the microbial community could be potential PAOs. Among them, only six genera had an average abundance > 0.5%, *Acinetobacter* was both the major potential PAO (median 21.5%) as well as the dominant genus (median 14.4%) during the period when the P-removal performance was good (Figure S4). Mehlig et al., (2013) investigated PAOs in eight WWTPs using combination of FACs and cloning, and found that *Acinetobacter* constituted about 20% of PAOs in a small oxidation ditch (500 m^3^ d^−1^). Eschenhagen et al., (2003) found 3 – 5 % *Acinetobacter* with *Tetrasphaera* as a major PAO (10%) in a lab-scale plant fed with municipal wastewater. In the present study, *Acinetobacter* abundance showed a significant positive correlation to P-release (*r* = 0.57, *p* < 0.05) but non-significant P-uptake rates (*r* = 0.34, *p* > 0.05) of periodic batch test (Table S4), indicating their potential involvement in P-removal. Thus, *Acinetobacter* activity may influence the P-removal performance considering their high relative abundance. *Tetrasphaera*, showed a weak correlation with P-release (r = 0.32, *p* > 0.05) and P-uptake rates (r = 0.49, *p* < 0.05), and its abundance is very low (median 0.05%), which did not support its major contribution to the P removal. No other candidate PAOs showed significant correlations with P removal.

### 3.4. Evidence of the involvement of *Acinetobacter* in EBPR

To further investigate the role and contribution of the Acinetobacter-like bacteria in our system to P removal, we first obtained and examined the P removal activity of the *Acinetobacter junni* (DSM14968) isolates from the DSMZ Cell Lines Bank (Braunschweig, Germany). It was found to have 98.8% similarity with the Acinetobacter-like OTU identified in our system (Figure S5). During the up-take and release test with this pure culture, both intracellular PHA and poly-P content increased continuously in anaerobic and aerobic phase (Figure S5D and S5E). In aerobic phase, the *A. junni* grows faster with acetate and thus utilize more PO_4_^3−^, but the intracellular poly-P grew slower than that without acetate (Figure S5E). Our results indicated that this *Acinetobacter strain* did not follow P release and uptake pattern in An/Aer cycle as for canonical Accumulibacter-like PAOs. Previous attempts also failed to confirm the cyclic phosphorus release and uptake ability of *Acinetobacter* that isolated from WWTP (Tandoi et al., 1998). Since the *Acinetobacter junni* strain may not represent the actual strain in our system, we therefore applied *in situ* phenotypic analysis by combing FISH and Raman spectroscopy techniques. Based on SCRS, cells featured with poly-P signals were considered as Raman identified PAOs. This approach could also be connected with a prior FISH step to target the intracellular poly-P content within *Acinetobacter spp*.

#### 3.4.1. Raman-FISH revealed the poly-P metabolism of *Acinetobacter-spp*

*Acinetobacter spp.* in from B-stage aerobic tank was identified by FISH-Raman analysis in an *ex-situ* An/Aer cycle experiment (Figure 2 and Table S4). The intracellular polyP content, as indicated by the obtained Raman signals intensities (13.82-546.4 counts), are comparable to our previously collected poly-P signal from *Ca*. Accumulibacter (5.39-1473.60 counts) and *Tetrasphaera* (6.78-1080.82 counts) using the similar SCRS acquisition conditions (Figure S8).

**Figure 2.**
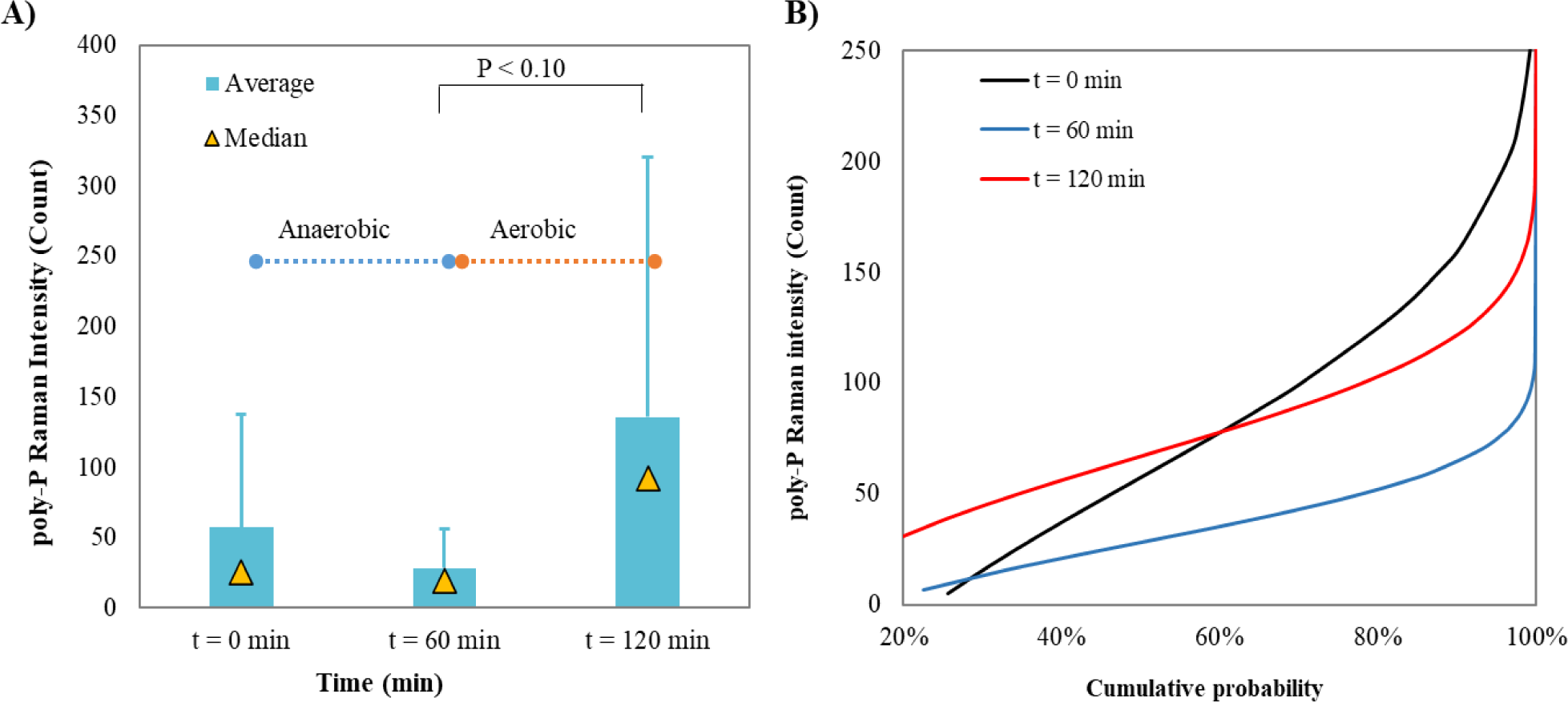
A) Polyphosphate Raman signal of *Acinetobacter* spp. and B) cumulative frequency distribution of poly-P signal in *ex-situ* P-release and P-uptake. t = 0, start of the test; t = 60 min, end of anaerobic / start of aerobic; t = 120, end of aerobic. Asterisks indicate marginally significant difference (P < 0.10).

The temporal changes of polyP content levels in *Acinetobacter* spp. based on the Raman-FISH signals revealed the potential polyp metabolism in *Acinetobacter*. Compared to the starting point of the anaerobic-aerobic cycle, the average poly-P signal of *Acinetobacter* spp. decreased by 51% at the end of the anaerobic (t = 60 min) and re-increased by 383% following the 60 min aeration (t = 120 min) (Figure 2A). The PO_4_^3−^ of the bulk solution showed an opposite trend to poly-P signal intensity (Figure S3).

This suggests that the *Acinetobacter spp*. in our HRAS-P(D)N-S2EBPR system seemed to display a PAO-like metabolism during the aerobic-anaerobic cycle. Further, the probability distribution of the poly-P signal was altered at each phase (Figure 2B), supporting the P-metabolism observations. However, the cumulative frequency of poly-P signals at the end of the test (t = 120 min) did not fully match the starting curve (t = 0 min), indicating that Acinetobacter spp. were actively accumulating poly-P, 60 min was insufficient for them to accumulate the poly-P level back to the beginning level.

#### 3.4.2. The involvement of *Acinetobacter*-like bacteria in the HRAS-P(D)N-S2EBPR system

In FISH-Raman procedure, fixation process is necessary for targeting the *Acinetobacter* spp. in sludge samples, but it may induce the loss/decrease of the intracellular polymer signals, including poly-P, PHA, and protein (Fernando et al., 2019). The loss was also observed in our preliminary study (data not shown). Thus, the poly-P signals for cells with low poly-P content could be completely lost after the fixation process, leading to underestimation of the relative abundance of PAOs and poly-P content levels. To overcome this obstacle, we applied the Raman-phenotypic profiling approach to identify the *in-situ* activity of *Acinetobacter*-like poly-P accumulating cells in the HRAS-P(D)N-S2EBPR system. Specifically, the fingerprint Raman signals from ACA23a labelled *Acinetobacter* spp. were obtained as a reference. All SCRS were obtained from the activated sludge and those closely aligned with the reference spectrum were considered *Acinetobacter*-like PAOs.

By this means, the changes of intracellular poly-P signals within all organisms and in *Acinetobacter*-like organisms (blue bars in Figure 3) can be obtained. The highest poly-P Raman signal in PAOs found in the S2EBPR was 2,933 counts, and this increased to 4,376 counts for the B-stage (Blue bars in Figure 3 and Table S5). The average poly-P signal of PAOs in the B-stage was marginally significantly higher (P < 0.10) than the S2EBPR.

**Figure 3.**
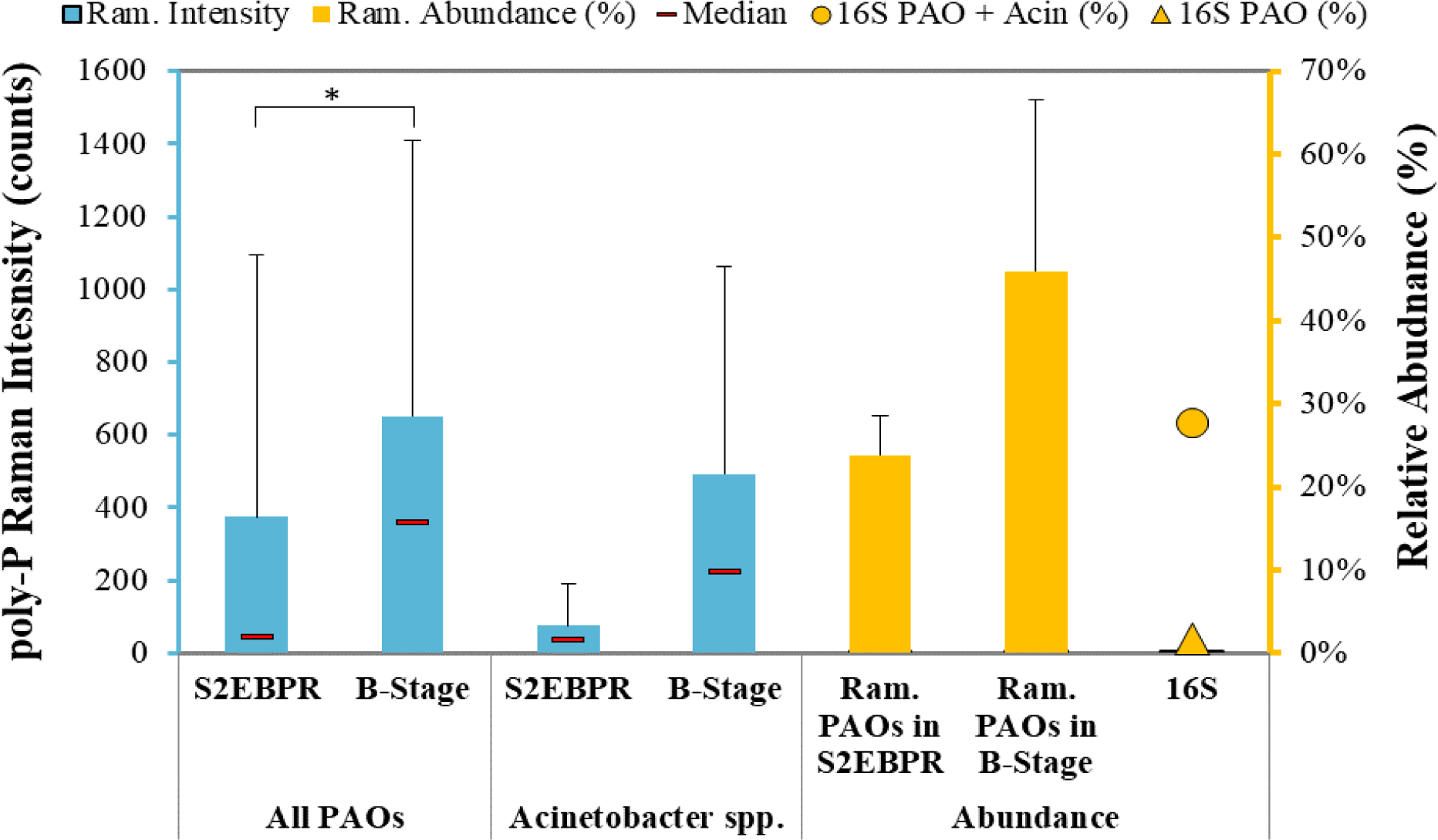
Average Poly-P signals in the poly-P containing cells identified by normal SCRS and *Acinetobacter* identified by FISH-SCRS (blue bars), and the SCRS-based phenotypic abundance (yellow bars), 16s-based phylogenic abundance of known PAOs. 16S PAO + Acin (%), abundance of genes related to known PAOs (Ca. Accumulibacter) and *Acinetobacter.* Asterisks indicate marginally significant difference (P < 0.10).

Based on SCRS phenotypic identification, the relative abundance of total PAOs among all organisms was obtained as 23.8 ± 4.7% in the S2EBPR, and this increased to 45.94 ± 20.7% by the end of the B-stage (yellow bars in Figure 3), evidencing the EBPR activities in this system. This relative PAOs abundance was significantly higher than the known PAOs identified by the 16S rRNA analysis (1.3 – 2.4%, circle in Figure 3), indicating the selection of unknow PAOs in this unique system. Categorizing *Acinetobacter* (26% at the time of the analysis from 16S sequencing) as a potential PAO can mitigate some gap, but the difference between the two was still substantial. Similarly, the total known GAO abundance identified using the genetic method via 16S rRNA sequencing was only 0.1 – 0.4%, which is a only a small fraction of those identified via SCRS with phenotypic features of GAO (ca. 6.9-24.1%). Thus, the 16S rRNA amplicon-based method may have some deficiency in identifying unknown yet active PAOs and GAOs in the HRAS-P(D)N-S2EBPR system (section 3.1.1). Nevertheless, the increase in the PAOs abundance and poly-P signal corroborates the PAO activity and net P-removal along the HRAS-P(D)N-S2EBPR train, suggesting enrichment of other non-caronical PAOs in this system.

*Acinetobacter*-like PAOs were ca. 54% of the total PAOs in the S2EBPR and the highest poly-P signals in the *Acinetobacter*-like PAOs is 356 count (Figure 3). A similar percentage of candidate PAOs in the B-stage were associated with *Acinetobacter*, and their highest poly-P signal was 2,289 counts. The intracellular poly-P signals indicated that *Acinetobacter*-like cells dominated the total PAOs population, and they accumulated more poly-P in B-stage aeration tank than in S2EBPR tank. The highest poly-P signal of *Acinetobacter*-like was lower than the highest value observed from total PAOs (Figure 3), suggesting that there are other PAOs that can store more poly-P per cell. Previous Raman-phenotypic profiling in a lab-scale EBPR system revealed that the cells with a very high poly-P signal are more closely related to the class β-Proteobacteria (Li et al., 2018), such as Ca. *Accumulibacter* - the most widely acknowledged PAO and *Dechloromonas* - a putative PAO. Nevertheless, the poly-P accumulating ability detected by both FISH-Raman and Raman phenotypic profiling and the dominant abundance of *Acinetobacter* all highly suggested its contribution to P removal in the HRAS-P(D)N-S2EBPR system.

#### 3.4.3. Diversity of Acinetobacter and Metabolic Capacity assessment

Conflicting results from pure culture and activated sludge indicated that there are diverse phenotypes of *Acinetobacter spp*., i.e., PAO-like and non-PAO like *Acinetobacter*. The micro diversity within the OTUs that identified as *Acinetobacter* were further resolved into oligotypes for assessment of their temporal oligotyping profiles (Figure 4). Based on the oligotyping, the most dominant OTU constituted ca. 97% *Acinetobacte-liker* OTU of the studied system. There were 20 oligotypes in the major OTU, among them the top seven oligos accounted for nearly 97% of the major OTU, with the remaining 13 oligotypes as minor composition. The two most abundant *Acinetobacter* oligos (ATTTAGA (1) and GTTTAGA (2)) were consistently dominant in the period of low effluent-P and good P removal. Linear regression indicated a negative correlation of effluent-P concentration with oligo1 (Figure S9, *r* = −0.58, *p* < 0.01), while no correlation with oligo2. This suggests that the phosphorus removal capacity may reside within the population of all *Acinetobacter* spp., specifically within oligo1.

**Figure 4.**
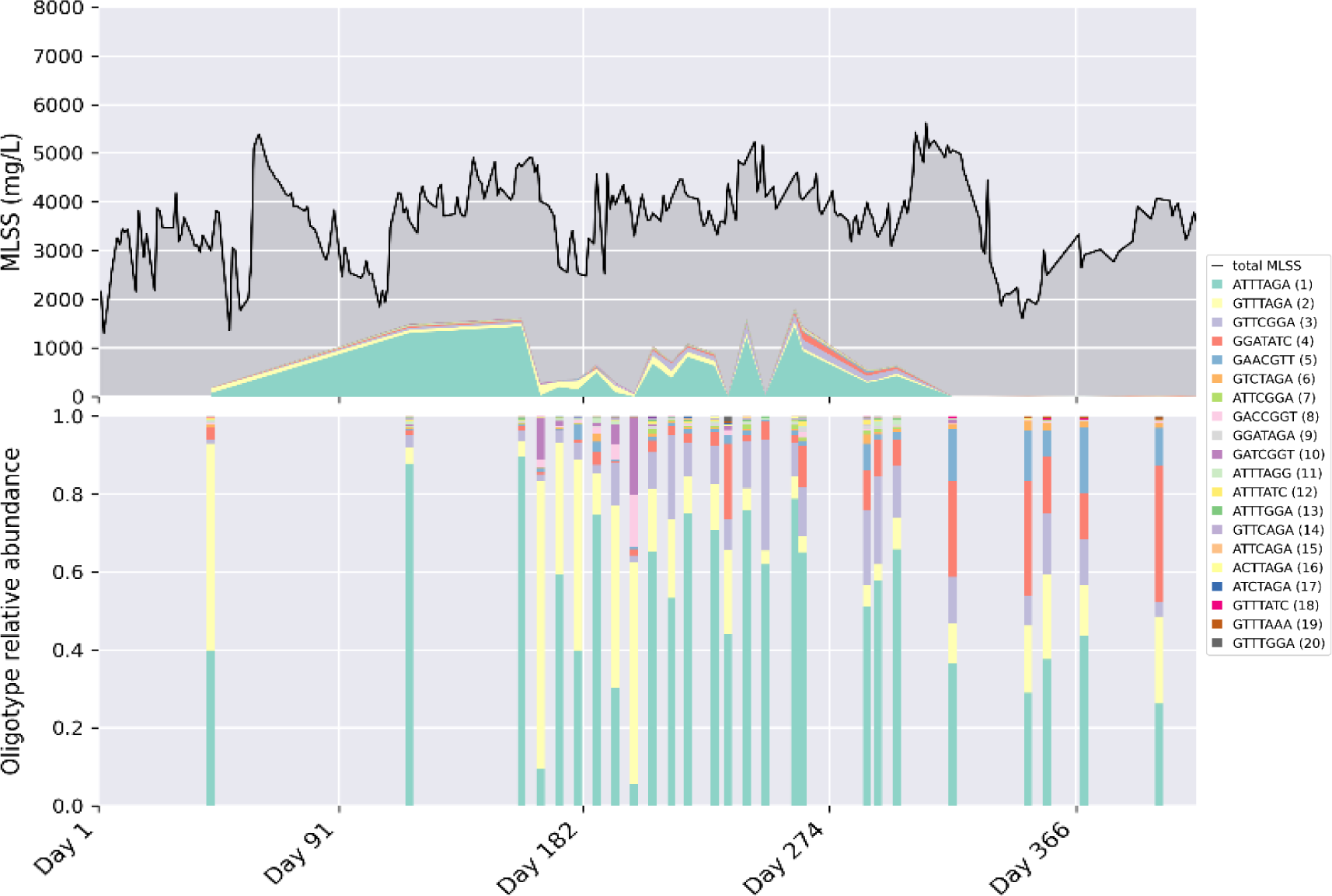
Relative abundance of oligotypes (20 total) of 16S rRNA gene amplicon sequences classified as genus *Acinetobacter* throughout the investigation period.

The 16S V4 of representative sequence for the oligotypes ATTTAGA (1) had identical (100% alignment identity) hits from *A. chinesis*, *A. gandensis*, and *A. bouvetti* in the nt database using BLAST, and the GTTAGA (2) had identical hit from *A. johnsonii*. Based on the literature review, *A. bouvetti* and *A. johnsonii* are phylogenetically close (Mateo-Estrada et al., 2019). Since metagenomic data of our sample was unavailable, core EBPR-related pathways of the identified species were analyzed using the assemblies downloaded from the NCBI assembly database to estimate or deduce their likely metabolic functions(Figure 5). Genetically these species can accumulate (*ppk1* and *ppk2*) and cleavage (*ppx*) poly-P. They can do partial denitrification NO_3_^−^ to NO (*nas* and *nir*), but not NO to N_2_, except *A. chinensis*. However, identified species may have different PHA accumulating metabolism from conventional PAOs, as they only seem to have the enzymes to make PHA monomers (*phaA* and *phaB*) as part of their metabolism.

**Figure 5.**
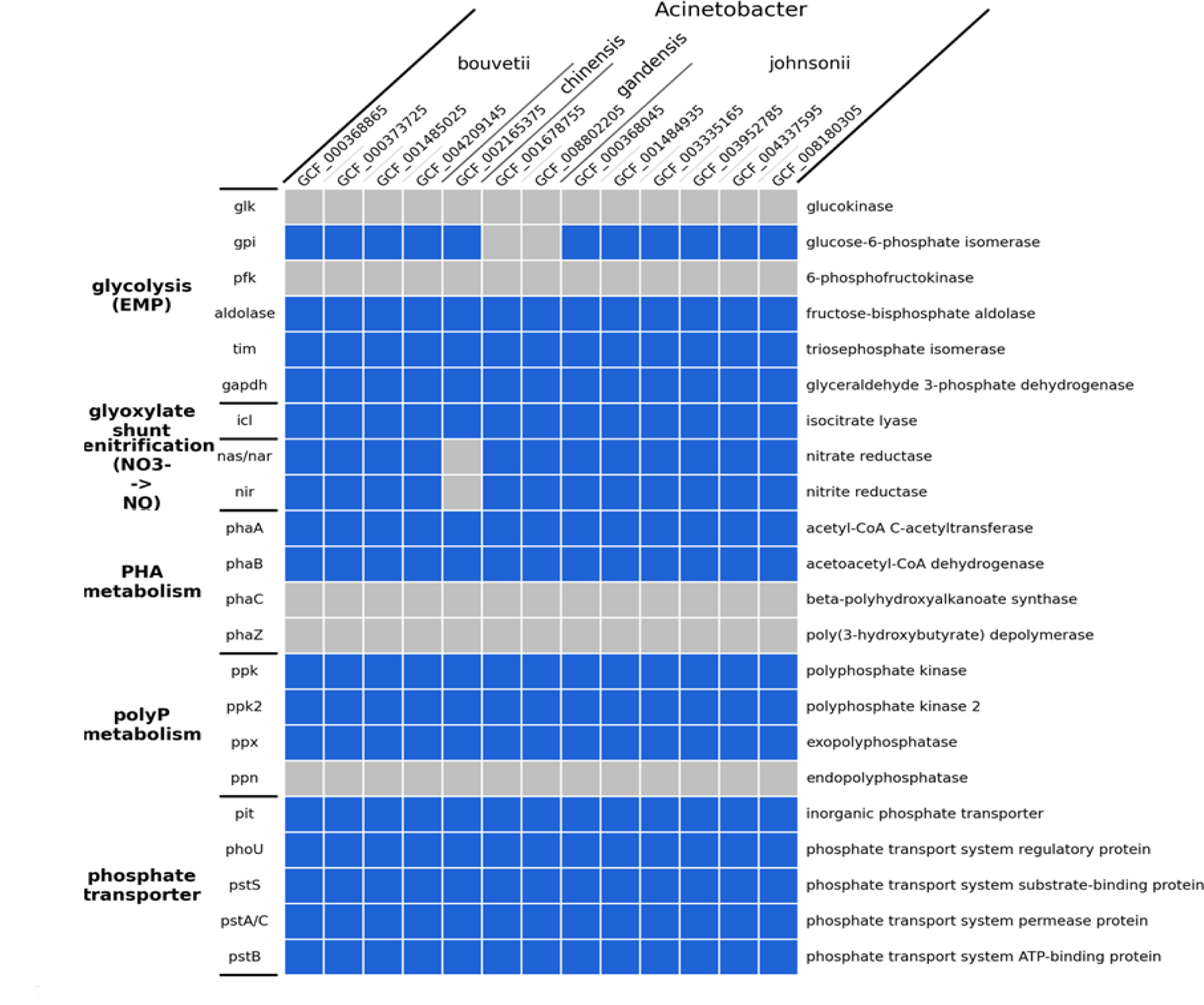
Genomic capabilities of *Acinetobacter* identified by oligotyping. A colored patch indicates a positive capability. The assessment was based on the annotated presence/absence of below essential genes in related pathways: fructose-bisphosphate aldolase; tim: triosephosphate isomerase; gapdh: glyceraldehyde 3-phosphate dehydrogenase; g6pd: glucose-6-phosphate 1-dehydrogenase; 6pgl: 6-phosphogluconolactonase; 2gp4: phosphogluconate dehydratase; kdpg aldolase: 2-dehydro-3-deoxyphosphogluconate aldolase; icl: isocitrate lyase; nas/nar: nitrate reductase; nir: nitrite reductase; nor: nitric oxide reductase; nos: nitrous-oxide reductase; nxr: nitrite oxidoreductase; amo: ammonia monooxygenase; hzo: hydroxylamine oxidase; gsy: glycogen synthase; glycogen phosphorylase; phaA: acetyl-CoA C-acetyltransferase; phaB: acetoacetyl-CoA dehydrogenase; phaC: poly(3-hydroxyalkanoate) polymerase; ppk: polyphosphate kinase; ppk2: polyphosphate kinase 2; ppx: exopolyphosphatase; pit: inorganic phosphate transporter; phoU: phosphate transport system regulatory protein; pstS: phosphate transport system substrate-binding protein; pstA/C: phosphate transport system permease protein; pstB: phosphate transport system ATP-binding protein.

## 4. Discussion

### 4.1. *Acinetobacter* contributed to phosphorus removal in HRAS-P(D)N-S2EBPR system

The role and contribution of *Acinetobacter* in full-scale EBPR performance has been a topic of discussion for decades. In this study,16S rRNA gene amplicon sequencing revealed the most dominant and surprisingly high relative abundance of Acinetobacter (16%), concurrently with very low relative abundance of known PAOs in our HRAS-P(D)N-S2EBPR system that exhibited efficient P removal. Previously, the major concern about the role of *Acinetobacter* in EBPR is raised because its P-uptake and P-release patterns does not correspond to conventional Accumulibacter-like PAOs in anaerobic/aerobic conditions (Tandoi et al., 1996). In the present study, Raman-phenotypic profiling and FISH-Raman analysis were employed to identify the Acinetobacter-like PAOs and quantify the intracellular poly-P signals at cellular and specific population resolution beyond conventional bulk methods. These results provided strong evidence that *Acinetobacter* is most likely involved in and contributed to the phosphorus removal in this unique system. There are over 30 species in the genus (Dijkshoorn and Nemec, 2008) and about five named species were reported to accumulate poly-P (Aravind et al., 2015; Hrenović et al., 2003; Ren et al., 2014; Zhang et al., 2019). Further microdiversity assessment via 16S sequencing coupled with oligotyping revealed two predominant *Acinetobacter* oligotypes in our system that had *ppk*/*ppx* in their genome for poly-P metabolism, and enzymes for PHA formation. These results pointed out the likely involvement and contribution of *Acinetobacter* in our unique HRAS-P(D)N-S2EBPR system that has distinct process configuration from the conventional EBPRs.

### 4.2. Versatile and metabolic traits of *Acinetobacter*

Overall, *Acinetobacter* have many unique metabolisms, such as aerobic/anoxic P-uptake with/without anaerobic stress (Liu et al., 2014); aerobic PHA accumulation (Preez et al., 1981; Zafiri et al., 1999); heterotrophic nitrification (Ren et al., 2014; Su et al., 2015) and resistance to antibiotics and pollutants (Margesin et al., 2003; Zhang et al., 2009). They can use diverse carbon sources and found in the primary sludge fermenter (Preez et al., 1981; Zafiri et al., 1999). Because of their versatile metabolism, *Acinetobacter* is resilient to environmental stress and is often found in contaminated environments. such as the sewerage system. *Acinetobacter* are non-fecal microorganisms, but they are one of the major genera found in the sewage water, showing that they can proliferate in sewerage pipes (McLellan and Roguet, 2019; VandeWalle et al., 2012). Interestingly, some of the conditions in sewerage pipeline fit relatively well with the HRAS-P(D)N-S2EBPR system. For instance, they all provide bacteria with aerobic/anaerobic condition. The total VFA concentration that flows into the mixing tank are around 13 – 60 mg-VFA L^−1^, which is close to that of the sewerage system (7 – 24 mg-VFA L^−1^) (Jin et al., 2015).

#### 4.2.1. Factors that may favor the proliferation of Acinetobacter

Despite the consistent dominance of *Acinetobacter* in this study, stable P-removal was observed only for a portion of the operation (Figure 4 and S4), coincided with increasing VFA/P ratio (Figure 1B, day 184 – 267). *Acinetobacter* possesses the genomic capacity to synthesize PHA (Figure S6) and were found from PHA accumulating mixed microbial cultures (MMC) (Dionisi et al., 2006; Villano et al., 2010; Zeng et al., 2018). Unlike pure culture, where glucose is mostly used as a substrate for PHA production, MMC biotechnology highly depend on the VFA as the precursors for PHA production (Kourmentza et al., 2017). By alternating the anaerobic-aerobic and anoxic-aerobic conditions, or by cycling the high/low organic loading rate (Dionisi et al., 2006), VFAs are stored by PHA accumulating microorganisms rapidly before OHOs can use them (Dionisi et al., 2006; Kourmentza et al., 2017). In the HRAS-P(D)N-S2EBPR system, S2EBPR creates 3.3 h anaerobic and VFA rich condition while the B-stage process creates 4.4-4.7 h anoxic/aerobic and VFA less condition, providing selective pressure on the activated sludge to accumulate PHA. Therefore, the rising mass flow of fermentate VFA to the S2EBPR (Figure 1B), could have given *Acinetobacter* an advantage over other non-carbon-accumulting OHOs in B-stage. The ability to accumulate PHA in VFA-rich S2EBPR can help them over compete OHOs when biodegradable COD is consumed.

The Raman-FISH and Raman phenotyping analysis provided molecular evidence that the P-removal performance of the HRAS-P(D)N-S2EBPR was likely attributed to the *Acinetobacter*-like PAOs that is the predominant organism during this period. However, the relative abundance did not correlate with the phosphorus removal performance directly. Other factors that may have influenced the performance are cyanide in the influent and NO_2_^−^ accumulation. Cyanide binds to Fe^3+^ of cytochrome oxidase system disrupting ATP production. The EBPR and *Acinetobacter* activity are inhibited by 0.10 mg-CN L^−1^ and 0.05 mg-CN L^−1^, respectively (Motlagh et al., 2015; Vierkant et al., 1990). The cyanide is an unlikely cause for performance deterioration as A-stage influent had an average cyanide of 0.014 ± 0.010 mg-CN L^−1^. Nitrite can inhibit PAO activity by increasing maintenance energy and disturbing the TCA cycle (Zhou et al., 2011). About 2 mg-NO_2_^−^-N L^−1^ at pH 7 can reduce the P-uptake rate of Ca. *Accumulibacter* by 50% (Pijuan et al., 2010). Similarly, 5 mg mg-NO_2_^−^-N L^−1^ at pH 6.6 can reduce the P-uptake rate of *A. junii* by 30% (Hrenović et al., 2003). Although a positive correlation was found between effluent PO_4_^3−^-P and NO_2_^−^-N (Figure 1A), it is unclear whether the increase in effluent-P is due to nitrite inhibition or installation of the pre-denitrification tank. Further investigation is required to identify the genus and mechanism responsible for NO_2_^−^ accumulation as well as optimal conditions to achieve simultaneous P removal and NO_2_^−^ accumulation.

## 5. key findings

1. Sufficient (85%) phosphorus removal was achieved (Eff. 0.63 ± 0.58 mg-P L^−1^) with low abundance of known PAOs (0.6 ± 0.3%) in a novel HRAS-P(D)N-S2EBPR system.
2. Combined 16s phylogenetic analysis, SCRS-based phenotypic profiling and FIHS-Raman metabolic tracing provided strong evidence that *Acinetobacter* most likely was the predominant PAO (median 16%) in our HRAS-P(D)N-S2EBPR system. Further, 16S rRNA-gene based amplicon sequencing and oligotyping analysis revealed subgroup of Acinetobacter (oligo1) might contribute to the phosphorus removal. Although previous studies excluded the role of *Acinetobacter* in EBPR due to its different metabolic features in terms of poly-P and PHA accumulation, our study pointed out the need to revisit its role in systems that has different configurations from conventional EBPRS.
3. To the best of our knowledge, this is the first study to confirm the involvement of *Acinetobacter* in EBPR using combined genomic and SCRS Raman techniques. Further research is needed to identify the specific taxon, and phenotype of the Acinetobacter that are responsible for the P-removal. To overcome this limitation, a better Raman signals identification method and Raman fingerprint database must be established so that conserved phenotypic properties can be identified under changing environmental conditions, and growth stages.

## Acknowledgement

This study was funded by the Water Environment Research Foundation (WERF, Grant No. 4901). We gratefully acknowledge the financial support and scientific discussion from Charles B. Bott (HRSD), Jim McQuarrie (Denver Metro), Beverley M. Stinson (AECOM), Christine deBarbadillo (DC Water), Paul Dombrowski (Woodard & Curran) and James Barnard (Black & Veatch). Special thanks are given to the generous support of operators and staff of HRSD for their assistance with sample and data collection.

## Reference

American Public Health Association, others, 2005. Standard methods for the examination of water and wastewater. APHA. American Water Works Association and Water Environment Federation, 21st ed.; American Public Health Association: Washington, DC, USA.

Aravind, J., Saranya, T., Kanmani, P., 2015. Optimizing the production of Polyphosphate from Acinetobacter towneri. Global Journal of Environmental Science and Management 1, 63–70.

Cole, J.R., Wang, Q., Fish, J.A., Chai, B., McGarrell, D.M., Sun, Y., Brown, C.T., Porras-Alfaro, A., Kuske, C.R., Tiedje, J.M., 2014. Ribosomal Database Project: data and tools for high throughput rRNA analysis. Nucl. Acids Res. 42, D633–D642.

Dionisi, D., Majone, M., Vallini, G., Di Gregorio, S., Beccari, M., 2006. Effect of the applied organic load rate on biodegradable polymer production by mixed microbial cultures in a sequencing batch reactor. Biotechnol. Bioeng. 93, 76–88.

Eren, A.M., Maignien, L., Sul, W.J., Murphy, L.G., Grim, S.L., Morrison, H.G., Sogin, M.L., 2013. Oligotyping: differentiating between closely related microbial taxa using 16S RRNA gene data. Methods Ecol Evol 4, 1111–1119.

Eschenhagen, M., Schuppler, M., Röske, I., 2003. Molecular characterization of the microbial community structure in two activated sludge systems for the advanced treatment of domestic effluents. Water Research 37, 3224–3232.

Fernando, E.Y., McIlroy, S.J., Nierychlo, M., Herbst, F.-A., Petriglieri, F., Schmid, M.C., Wagner, M., Nielsen, J.L., Nielsen, P.H., 2019. Resolving the individual contribution of key microbial populations to enhanced biological phosphorus removal with Raman– FISH. ISME J 13, 1933–1946.

Filipe, C.D.M., Daigger, G.T., Grady, C.P.L., 2001. pH as a Key Factor in the Competition Between Glycogen-Accumulating Organisms and Phosphorus-Accumulating Organisms. Water Environment Research 73, 223–232.

Fuhs, G.W., Chen, M., 1975. Microbiological basis of phosphate removal in the activated sludge process for the treatment of wastewater. Microb Ecol 2, 119–138.

Gu, A.Z., Saunders, A., Neethling, J.B., Stensel, H.D., Blackall, L.L., 2008. Functionally Relevant Microorganisms to Enhanced Biological Phosphorus Removal Performance at Full-Scale Wastewater Treatment Plants in the United States. Water Environment Research 80, 688–698.

Gu, A.Z., Tooker, N.B., Onnis-Hayden, A., Wang, D., Li, G., Srinivasan, V., Takács, I., 2018. Investigation of the Mechanisms for Optimization and Design of a Side-Stream EBPR Process as a Sustainable Approach for Achieving Stable and Efficient Phosphorus Removal. by Water Environmental Research Foundation (WERF) and International Water Association (IWA) Publishing.

Han, Y.-H., Fu, T., Wang, S.-S., Yu, H.-T., Xiang, P., Zhang, W.-X., Chen, D.-L., Li, M., 2018. Efficient phosphate accumulation in the newly isolated Acinetobacter junii strain LH4. 3 Biotech 8, 313.

Hrenović, J., Tibljaš, D., Büyükgüngör, H., Orhan, Y., 2003. Influence of Support Materials on Phosphate Removal by the Pure Culture of Acinetobacter calcoaceticus. Food Technology and Biotechnology 41, 331–338.

Hyatt, D., Chen, G.-L., LoCascio, P.F., Land, M.L., Larimer, F.W., Hauser, L.J., 2010. Prodigal: prokaryotic gene recognition and translation initiation site identification. BMC Bioinformatics 11, 119.

Jin, P., Wang, Bin, Jiao, D., Sun, G., Wang, Baobao, Wang, X.C., 2015. Characterization of microflora and transformation of organic matters in urban sewer system. Water Research 84, 112–119.

Kanehisa, M., 2006. From genomics to chemical genomics: new developments in KEGG. Nucleic Acids Research 34, D354–D357.

Kanehisa, M., Sato, Y., Morishima, K., 2016. BlastKOALA and GhostKOALA: KEGG Tools for Functional Characterization of Genome and Metagenome Sequences. Journal of Molecular Biology 428, 726–731.

Kang, D., Han, I.L., Lee, J., McCullough, K., Li, G., Wang, D., Klaus, S., Zheng, P., Srinivasan, V., Bott, C., Gu, A.Z., 2023. Molecular Evidence of Internal Carbon-Driven Partial Denitrification Annamox (PdNA) in a mainstream Pilot A-B System Coupled with Side-stream EBPR treating municipal wastewater.

Kawakoshi, A., Nakazawa, H., Fukada, J., Sasagawa, M., Katano, Y., Nakamura, S., Hosoyama, A., Sasaki, H., Ichikawa, N., Hanada, S., Kamagata, Y., Nakamura, K., Yamazaki, S., Fujita, N., 2012. Deciphering the Genome of Polyphosphate Accumulating Actinobacterium Microlunatus phosphovorus. DNA Research 19, 383–394.

Klaus, S., Printz, K., McCullough, K., Srinivasan, V., Wang, D., He, P., De Clippeleir, H., Gu, A., Bott, C., 2019. Integrating BioP and Shortcut Nitrogen Removal via RAS Fermentation and Partial Denitrification/Anammox. Presented at the Nutrient Removal and Recovery Symposium 2019, Water Environment Federation.

Klaus, S.A., 2019. Intensification of Biological Nutrient Removal Processes (Doctor of Philosophy). Virginia Polytechnic Institute and State University, Blacksburt, Virginia.

Kourmentza, C., Plácido, J., Venetsaneas, N., Burniol-Figols, A., Varrone, C., Gavala, H.N., Reis, M.A.M., 2017. Recent Advances and Challenges towards Sustainable Polyhydroxyalkanoate (PHA) Production. Bioengineering 4, 55.

Kristiansen, R., Nguyen, H.T.T., Saunders, A.M., Nielsen, J.L., Wimmer, R., Le, V.Q., McIlroy, S.J., Petrovski, S., Seviour, R.J., Calteau, A., Nielsen, K.L., Nielsen, P.H., 2013. A metabolic model for members of the genus Tetrasphaera involved in enhanced biological phosphorus removal. ISME J 7, 543–554.

Lackner, S., Horn, H., 2013. Comparing the performance and operation stability of an SBR and MBBR for single-stage nitritation-anammox treating wastewater with high organic load. Environmental Technology 34, 1319–1328.

Li, G., Wu, C., Wang, D., Srinivasan, V., Kaeli, D.R., Dy, J.G., Gu, A.Z., 2022. Machine Learning-Based Determination of Sampling Depth for Complex Environmental Systems: Case Study with Single-Cell Raman Spectroscopy Data in EBPR Systems. Environ. Sci. Technol. 56, 13473–13484.

Li, X., Yuan, H., Yang, J., Li, B., 2013. Genome Sequence of the Polyphosphate-Accumulating Organism *Arthrobacter* sp. Strain PAO19 Isolated from Maize Rhizosphere Soil. Genome Announc 1, e00566-13.

Li, Y., Cope, H.A., Rahman, S.M., Li, G., Nielsen, P.H., Elfick, A., Gu, A.Z., 2018. Toward Better Understanding of EBPR Systems via Linking Raman-Based Phenotypic Profiling with Phylogenetic Diversity. Environ. Sci. Technol. 52, 8596–8606.

Liu, Y., Li, X., Kang, X., Yuan, Y., 2014. Performance of denitrifying phosphorus removal of Acinetobacteria strain at low temperature. International Biodeterioration & Biodegradation, Challenges in Environmental Science and Engineering, CESE-2013 95, 135–138.

Lopez-Vazquez, C.M., Oehmen, A., Hooijmans, C.M., Brdjanovic, D., Gijzen, H.J., Yuan, Z., van Loosdrecht, M.C.M., 2009. Modeling the PAO–GAO competition: Effects of carbon source, pH and temperature. Water Research 43, 450–462.

Majed, N., Gu, A.Z., 2010. Application of Raman Microscopy for Simultaneous and Quantitative Evaluation of Multiple Intracellular Polymers Dynamics Functionally Relevant to Enhanced Biological Phosphorus Removal Processes. Environ. Sci. Technol. 44, 8601–8608.

Margesin, R., Labbé, D., Schinner, F., Greer, C.W., Whyte, L.G., 2003. Characterization of Hydrocarbon-Degrading Microbial Populations in Contaminated and Pristine Alpine Soils. Appl Environ Microbiol 69, 3085–3092.

Martín, H.G., Ivanova, N., Kunin, V., Warnecke, F., Barry, K.W., McHardy, A.C., Yeates, C., He, S., Salamov, A.A., Szeto, E., Dalin, E., Putnam, N.H., Shapiro, H.J., Pangilinan, J.L., Rigoutsos, I., Kyrpides, N.C., Blackall, L.L., McMahon, K.D., Hugenholtz, P., 2006. Metagenomic analysis of two enhanced biological phosphorus removal (EBPR) sludge communities. Nat Biotechnol 24, 1263–1269.

Mateo-Estrada, V., Graña-Miraglia, L., López-Leal, G., Castillo-Ramírez, S., 2019. Phylogenomics Reveals Clear Cases of Misclassification and Genus-Wide Phylogenetic Markers for Acinetobacter. Genome Biology and Evolution 11, 2531– 2541.

McCullough, K., Klaus, S., Schoepflin, S., Wilson, C., Gu, A., Han, I., Clippeleir, H. de, Bott, C., 2020. Simultaneous N and P Removal with Anammox and Sidestream BioP. Presented at the WEFTEC 2020, Water Environment Federation.

McLellan, S.L., Roguet, A., 2019. The unexpected habitat in sewer pipes for the propagation of microbial communities and their imprint on urban waters. Current Opinion in Biotechnology 57, 34–41.

Mehlig, L., Petzold, M., Heder, C., Günther, S., Müller, S., Eschenhagen, M., Röske, I., Röske, K., 2013. Biodiversity of Polyphosphate Accumulating Bacteria in Eight WWTPs with Different Modes of Operation. J. Environ. Eng. 139, 1089–1098.

Miller, M.W., Elliott, M., DeArmond, J., Kinyua, M., Wett, B., Murthy, S., Bott, C.B., 2017. Controlling the COD removal of an A-stage pilot study with instrumentation and automatic process control. Water Science and Technology 75, 2669–2679.

Motlagh, A.M., Bhattacharjee, A.S., Goel, R., 2015. Microbiological study of bacteriophage induction in the presence of chemical stress factors in enhanced biological phosphorus removal (EBPR). Water Research 81, 1–14.

Neethling, J.B., Bakke, B., Benisch, M., Gu, A.Z., Stephens, H., Stensel, H.D., Moore, R., 2006. Factors Influencing the Reliability of Enhanced Biological Phosphorus Removal (Final report No. 01-CTS-3). Alexandria, VA: Water Environment Research Foundation.

Oehmen, A., Yuan, Z., Blackall, L.L., Keller, J., 2005. Comparison of acetate and propionate uptake by polyphosphate accumulating organisms and glycogen accumulating organisms. Biotechnology and Bioengineering 91, 162–168.

Onnis-Hayden, A., Srinivasan, V., Tooker, N.B., Li, G., Wang, D., Barnard, J.L., Bott, C., Dombrowski, P., Schauer, P., Menniti, A., Shaw, A., Stinson, B., Stevens, G., Dunlap, P., Takács, I., McQuarrie, J., Phillips, H., Lambrecht, A., Analla, H., Russell, A., Gu, A.Z., 2020. Survey of full-scale sidestream enhanced biological phosphorus removal (S2EBPR) systems and comparison with conventional EBPRs in North America: Process stability, kinetics, and microbial populations. Water Environment Research 92, 403–417.

Pijuan, M., Ye, L., Yuan, Z., 2010. Free nitrous acid inhibition on the aerobic metabolism of poly-phosphate accumulating organisms. Water Research 44, 6063–6072.

Preez, J.C., Toerien, D.F., Lategan, P.M., 1981. Growth parameters of Acinetobacter calcoaceticus on acetate and ethanol. European J. Appl. Microbiol. Biotechnol. 13, 45– 53.

Quast, C., Pruesse, E., Yilmaz, P., Gerken, J., Schweer, T., Yarza, P., Peplies, J., Glöckner, F.O., 2012. The SILVA ribosomal RNA gene database project: improved data processing and web-based tools. Nucleic Acids Research 41, D590–D596.

R Core Team, 2013. R: A Language and Environment for Statistical Computing. R Foundation for Statistical Computing, Vienna, Austria.

Ren, Y.-X., Yang, L., Liang, X., 2014. The characteristics of a novel heterotrophic nitrifying and aerobic denitrifying bacterium, Acinetobacter junii YB. Bioresource Technology 171, 1–9.

Saunders, A.M., Albertsen, M., Vollertsen, J., Nielsen, P.H., 2016. The activated sludge ecosystem contains a core community of abundant organisms. ISME J 10, 11–20.

Schloss, P.D., Westcott, S.L., Ryabin, T., Hall, J.R., Hartmann, M., Hollister, E.B., Lesniewski, R.A., Oakley, B.B., Parks, D.H., Robinson, C.J., Sahl, J.W., Stres, B., Thallinger, G.G., Van Horn, D.J., Weber, C.F., 2009. Introducing mothur: Open-Source, Platform-Independent, Community-Supported Software for Describing and Comparing Microbial Communities. Appl Environ Microbiol 75, 7537–7541.

Soo, R.M., Skennerton, C.T., Sekiguchi, Y., Imelfort, M., Paech, S.J., Dennis, P.G., Steen, J.A., Parks, D.H., Tyson, G.W., Hugenholtz, P., 2014. An Expanded Genomic Representation of the Phylum Cyanobacteria. Genome Biology and Evolution 6, 1031–1045.

Stokholm-Bjerregaard, M., McIlroy, S.J., Nierychlo, M., Karst, S.M., Albertsen, M., Nielsen, P.H., 2017. A Critical Assessment of the Microorganisms Proposed to be Important to Enhanced Biological Phosphorus Removal in Full-Scale Wastewater Treatment Systems. Frontiers in Microbiology 8.

Su, J., Zhang, K., Huang, T., Wen, G., Guo, L., Yang, S., 2015. Heterotrophic nitrification and aerobic denitrification at low nutrient conditions by a newly isolated bacterium, Acinetobacter sp. SYF26. Microbiology 161, 829–837.

Tandoi, V., Majone, M., May, J., Ramadori, R., 1998. The behaviour of polyphosphate accumulating acinetobacter isolates in an anaerobic–aerobic chemostat. Water Research 32, 2903–2912.

VandeWalle, J.L., Goetz, G.W., Huse, S.M., Morrison, H.G., Sogin, M.L., Hoffmann, R.G., Yan, K., McLellan, S.L., 2012. Acinetobacter, Aeromonas and Trichococcus populations dominate the microbial community within urban sewer infrastructure: Dominant microbial populations of sewer infrastructure. Environmental Microbiology 14, 2538–2552.

Vierkant, M.A., Martin, D.W., Stewart, J.R., 1990. Poly-β-hydroxybutyrate production in eight strains of the genus *Acinetobacter*. Can. J. Microbiol. 36, 657–663.

Villano, M., Beccari, M., Dionisi, D., Lampis, S., Miccheli, A., Vallini, G., Majone, M., 2010. Effect of pH on the production of bacterial polyhydroxyalkanoates by mixed cultures enriched under periodic feeding. Process Biochemistry 45, 714–723.

Wagner, M., Erhart, R., Manz, W., Amann, R., Lemmer, H., Wedi, D., Schleifer, K.H., 1994. Development of an rRNA-targeted oligonucleotide probe specific for the genus Acinetobacter and its application for in situ monitoring in activated sludge. Appl Environ Microbiol 60, 792–800.

Wang, D., Tooker, N.B., Srinivasan, V., Li, G., Fernandez, L.A., Schauer, P., Menniti, A., Maher, C., Bott, C.B., Dombrowski, P., Barnard, J.L., Onnis-Hayden, A., Gu, A.Z., 2019. Side-stream enhanced biological phosphorus removal (S2EBPR) process improves system performance - A full-scale comparative study. Water Research 167, 115109.

Wang, X., Yang, R., Guo, Y., Zhang, Z., Kao, C.M., Chen, S., 2019. Investigation of COD and COD/N ratio for the dominance of anammox pathway for nitrogen removal via isotope labelling technique and the relevant bacteria. Journal of Hazardous Materials 366, 606– 614.

Wu, L., Ning, D., Zhang, B., Li, Y., Zhang, P., Shan, X., Zhang, Q., Brown, M.R., Li, Z., Van Nostrand, J.D., Ling, F., Xiao, N., Zhang, Y., Vierheilig, J., Wells, G.F., Yang, Y., Deng, Y., Tu, Q., Wang, A., Zhang, T., He, Z., Keller, J., Nielsen, P.H., Alvarez, P.J.J., Criddle, C.S., Wagner, M., Tiedje, J.M., He, Q., Curtis, T.P., Stahl, D.A., Alvarez-Cohen, L., Rittmann, B.E., Wen, X., Zhou, J., 2019. Global diversity and biogeography of bacterial communities in wastewater treatment plants. Nat Microbiol 4, 1183–1195.

Zafiri, C., Kornaros, M., Lyberatos, G., 1999. Kinetic modelling of biological phosphorus removal with a pure culture of Acinetobacter sp. under aerobic, anaerobic and transient operating conditions. Water Research 33, 2769–2788.

Zeng, S., Song, F., Lu, P., He, Q., Zhang, D., 2018. Improving PHA production in a SBR of coupling PHA-storing microorganism enrichment and PHA accumulation by feed-on-demand control. AMB Expr 8, 97.

Zhang, W., Gong, J., Wu, S., Yin, H., Jin, Y., Wu, H., Li, P., Wang, R., 2019. Draft genome Sequence of Phosphate-Accumulating Bacterium Acinetobacter tandoii SC36 from a Mangrove Wetland Ecosystem Provides Insights into Elements of Phosphorus Removal. Curr Microbiol 76, 207–212.

Zhang, Y., Marrs, C.F., Simon, C., Xi, C., 2009. Wastewater treatment contributes to selective increase of antibiotic resistance among Acinetobacter spp. Science of The Total Environment 407, 3702–3706.

Zhang, Z., Zhang, Y., Chen, Y., 2020. Recent advances in partial denitrification in biological nitrogen removal: From enrichment to application. Bioresource Technology 298, 122444.

Zhou, Y., Oehmen, A., Lim, M., Vadivelu, V., Ng, W.J., 2011. The role of nitrite and free nitrous acid (FNA) in wastewater treatment plants. Water Research 45, 4672–4682.

