## Supplementary material for "Revisiting the role of *Acinetobacter* spp. in EBPR systems": Suplemental Information

**Contents:**

**Figures**

Figure S1 Configuration of the P(D)N-S2EBPR pilot plant operating at HRSD Chesapeake-Elizabeth Plant in Virginia Beach, VA.	5

Figure S2 Cumulative-relative frequency distribution of efﬂuent PO_4_^-^-P of the P(D)N-S2EBPR system. Total, all efﬂuent PO_4_^-^-P measurements taken in the study. Colored dotted lines, exponential trend line of respective periods.	6

Figure S3 PAO activity profile taken on day 226, showing phosphorus release/uptake and intracellular polymer concentration variation during a anaerobic/aerobic cycle.	7

Figure S4 Non-metric multidimensional scaling (NMDS) plot of the Bray-Curtis similarity between bacterial community of the activated sludge at different time points. Numbers indicate sampling date. Blue numbers, before / after S2EBPR tank setup (before day 0), period of optimization period (days 1 – 147), and period of high effluent PO_4_^-^-P (days 268 – 355). Green number, after change in RAS% (days 150 – 267). Red number, period of effluent PO_4_^-^-P < 1.0 mg-P L^-1^ (days 184 – 244).	8

Figure S5 Relative abundance changes of (core) microorganisms found in the P(D)N-S2EBPR system of this study.	9

Figure S6 The variation of A) OD600, B) acetate, and C) PO_4_^3–^ concentration in the medium, and the normalized D) intracellular PHA and E) polyp intensity in individual cells in Anaerobic/Aerobic cycle analysis of pure-cultured *Acinetobacter junii*. Grey shadows indicate anaerobic phases.	10

Figure S7 Average PHA signals in the polyP containing cells identified by normal SCRS and in the *Acinetobacter* identified by FISH-SCRS (orange bars), and the SCRS-based phenotypic abundance (green bars).	11

Figure S8 Average PolyP signals in *Acinetobacter*, *Accumulibacter* and *Tetrasphaera* identified by FISH-SCRS (blue bars),	12

Figure S9 Linear correlation between the relative abundance of oligo1 (ATTTAGA) and the effluent-P concentrations (mg-P/L)	13

5 Tables

Table S 1. Influent and effluent PO_4_^-^-P of the P(D)N-S2EBPR system, and other full-scale EBPR and S2EBPR facilities. (Unit:)	14

Table S 2. *ex-situ* P-uptake and P-release tests for the P(D)N-S2EBPR system and other full-scale EBPR and S2EBPR facilities.	16

Table S 3 Diversity indices of 16S rRNA gene amplicon of the P(D)N-S2EBPR system in different periods	18

Table S4. Regression analysis of the abundance of *Acinetobacter* and *Tetrasphera* (%) to the anaerobic phosphorus release rate and aerobic phosphorus uptake rate (mg-P g_VSS_^-1^ h^-1^).	19

Table S 5. Summary of poly-P Raman signal of samples from the SBPR, B-stage reactor and FISH-Raman analysis of *ex-situ* P-release and P-uptake test. (Units: Count)	20

1 Text

Pure culture analysis of *Acinetobacter junni*	20

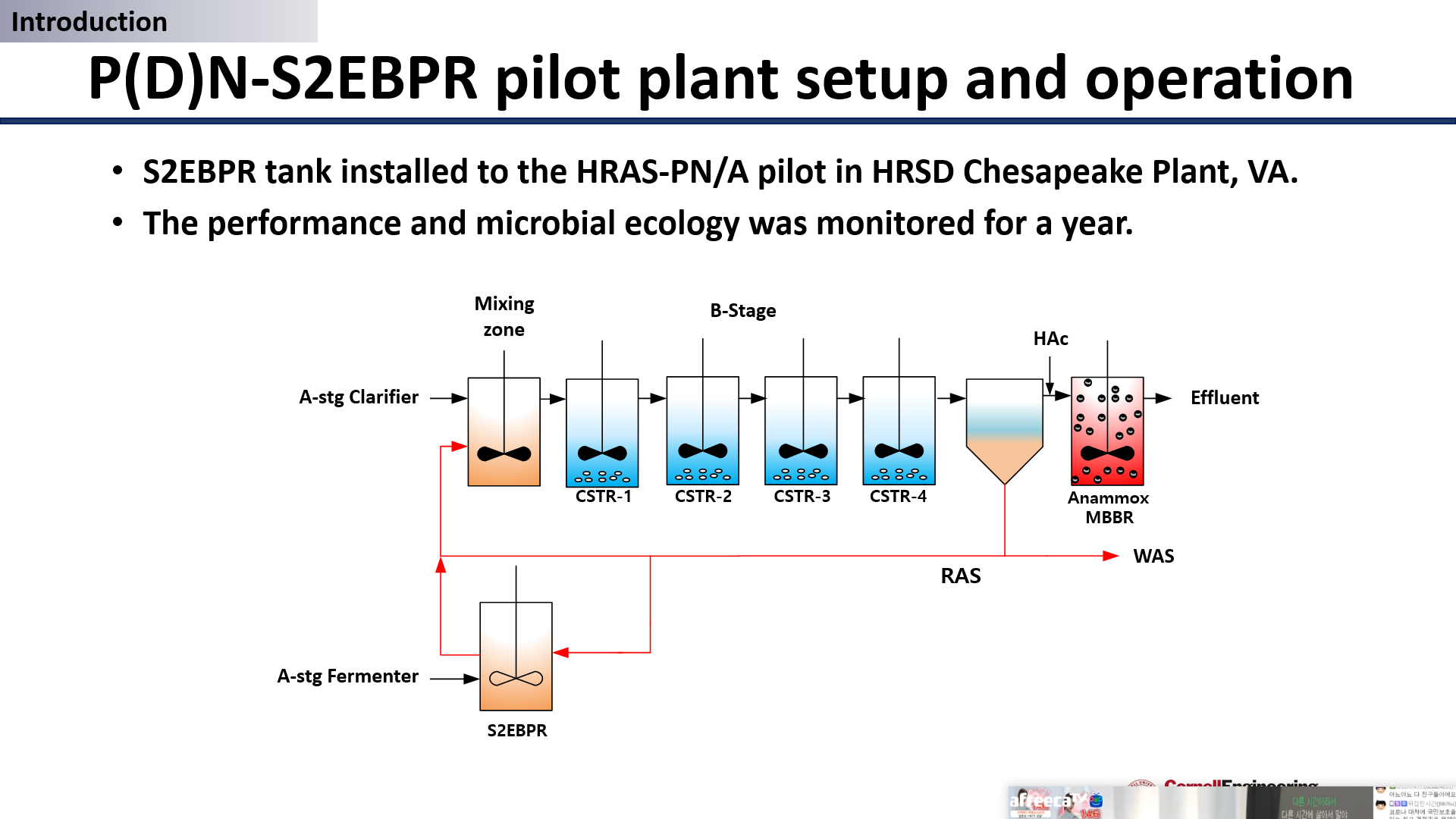

Figure S1 Configuration of the P(D)N-S2EBPR pilot plant operating at HRSD Chesapeake-Elizabeth Plant in Virginia Beach, VA.

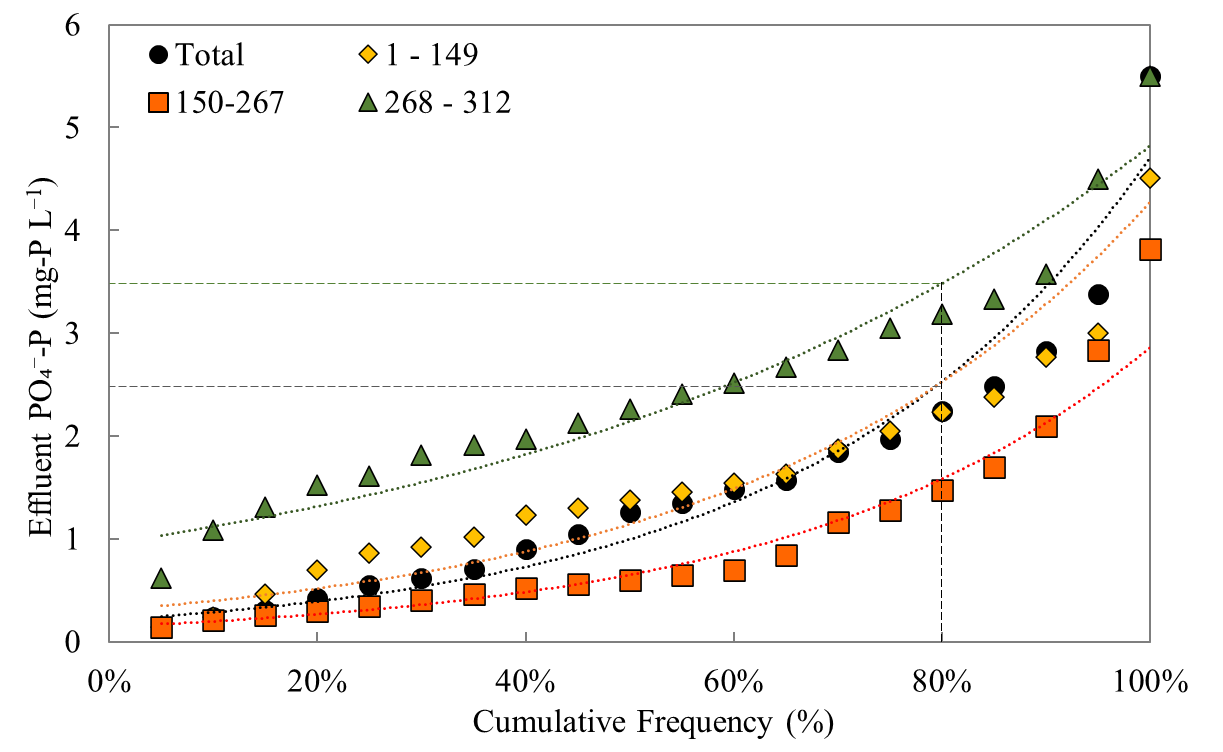

Figure S2 Cumulative-relative frequency distribution of efﬂuent PO_4_^-^-P of the P(D)N-S2EBPR system. Total, all efﬂuent PO_4_^-^-P measurements taken in the study. Colored dotted lines, exponential trend line of respective periods.

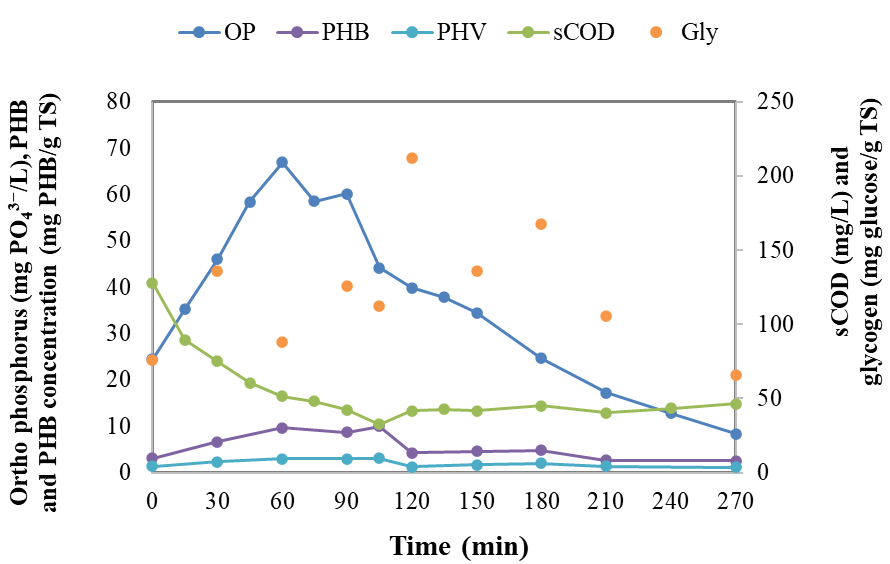

Figure S3 PAO activity profile taken on day 226, showing phosphorus release/uptake and intracellular polymer concentration variation during an anaerobic/aerobic cycle.

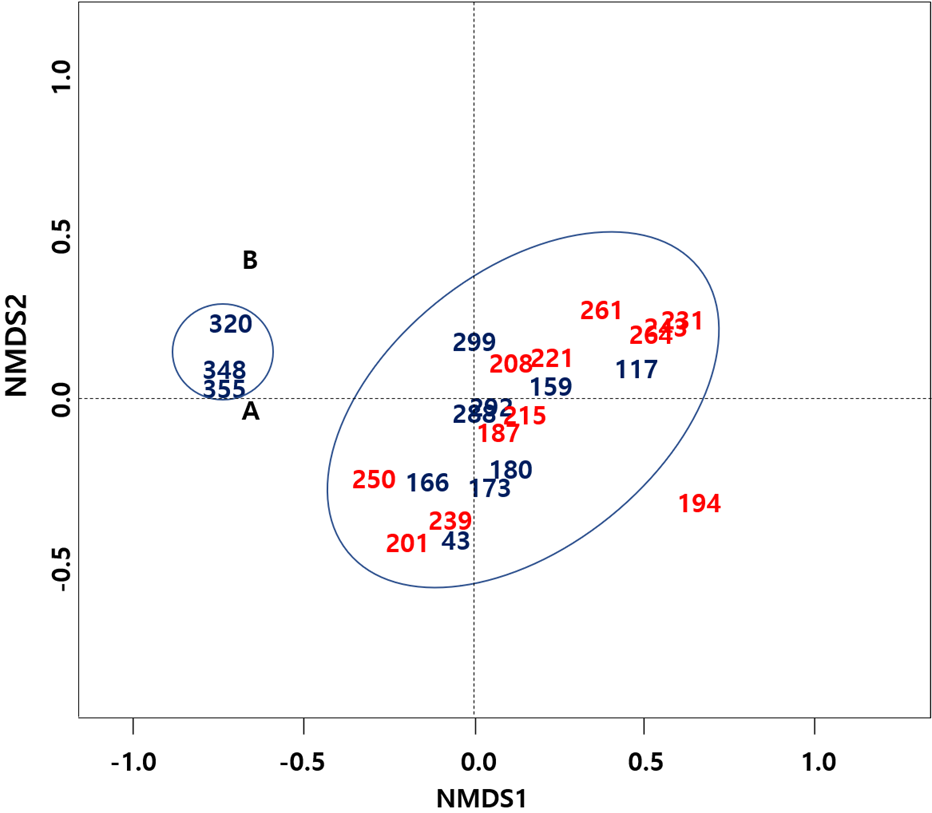

Figure S4 Non-metric multidimensional scaling (NMDS) plot of the Bray-Curtis similarity between bacterial community of the activated sludge at different time points. Numbers indicate sampling date. Blue numbers, before / after S2EBPR tank setup (before day 0), period of optimization period (days 1 – 147), and period of high effluent PO_4_^-^-P (days 268 – 355). Green number, after a change in RAS% (days 150 – 267). Red number, period of effluent PO_4_^-^-P < 1.0 mg-P L^-1^ (days 184 – 244).

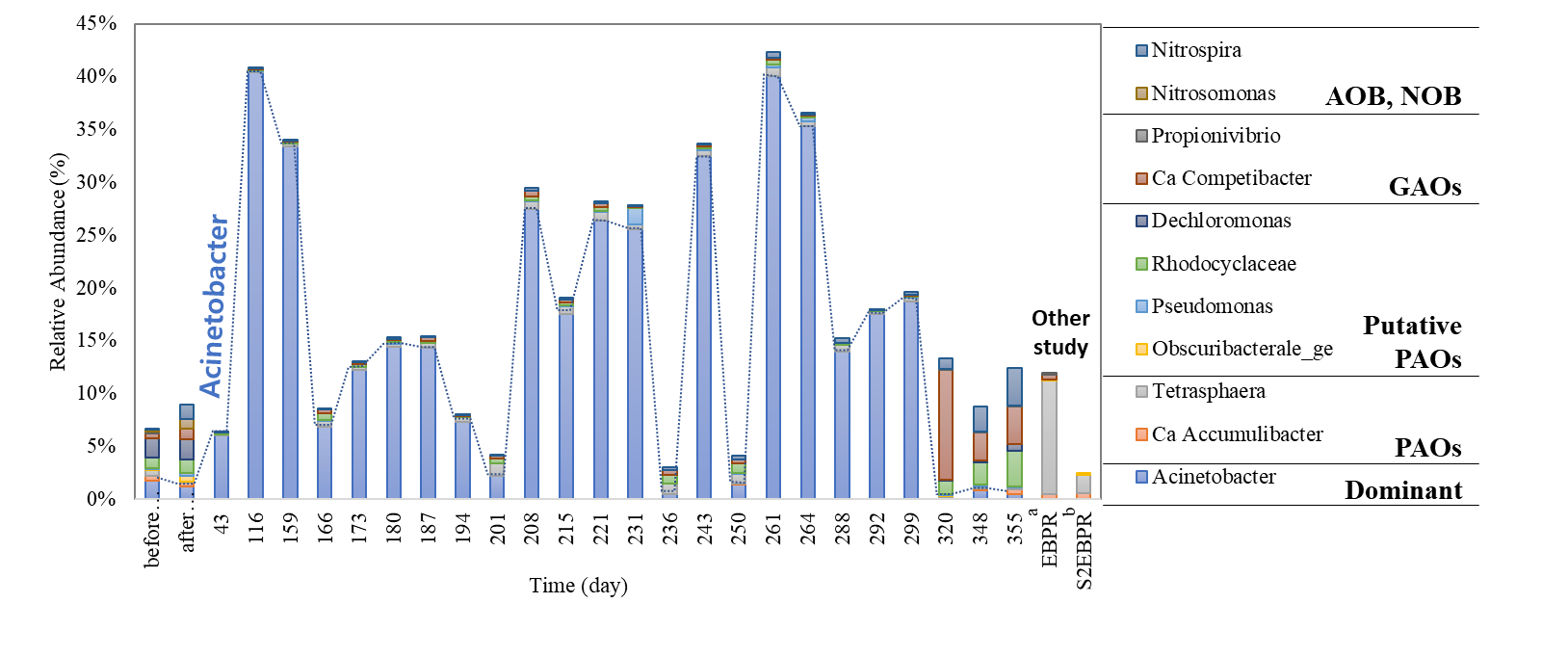

Figure S5 Relative abundance changes of (core) microorganisms found in the P(D)N-S2EBPR system of this study.

The list includes dominant genus, conventional PAOs, GAOs, putative PAOs, AOB and NOBs found in the 16S rRNA amplicon analysis. The abundances were compared with microbial communities from other studies including a: (Onnis-Hayden et al., 2020) and b: (Stokholm-Bjerregaard et al., 2017)

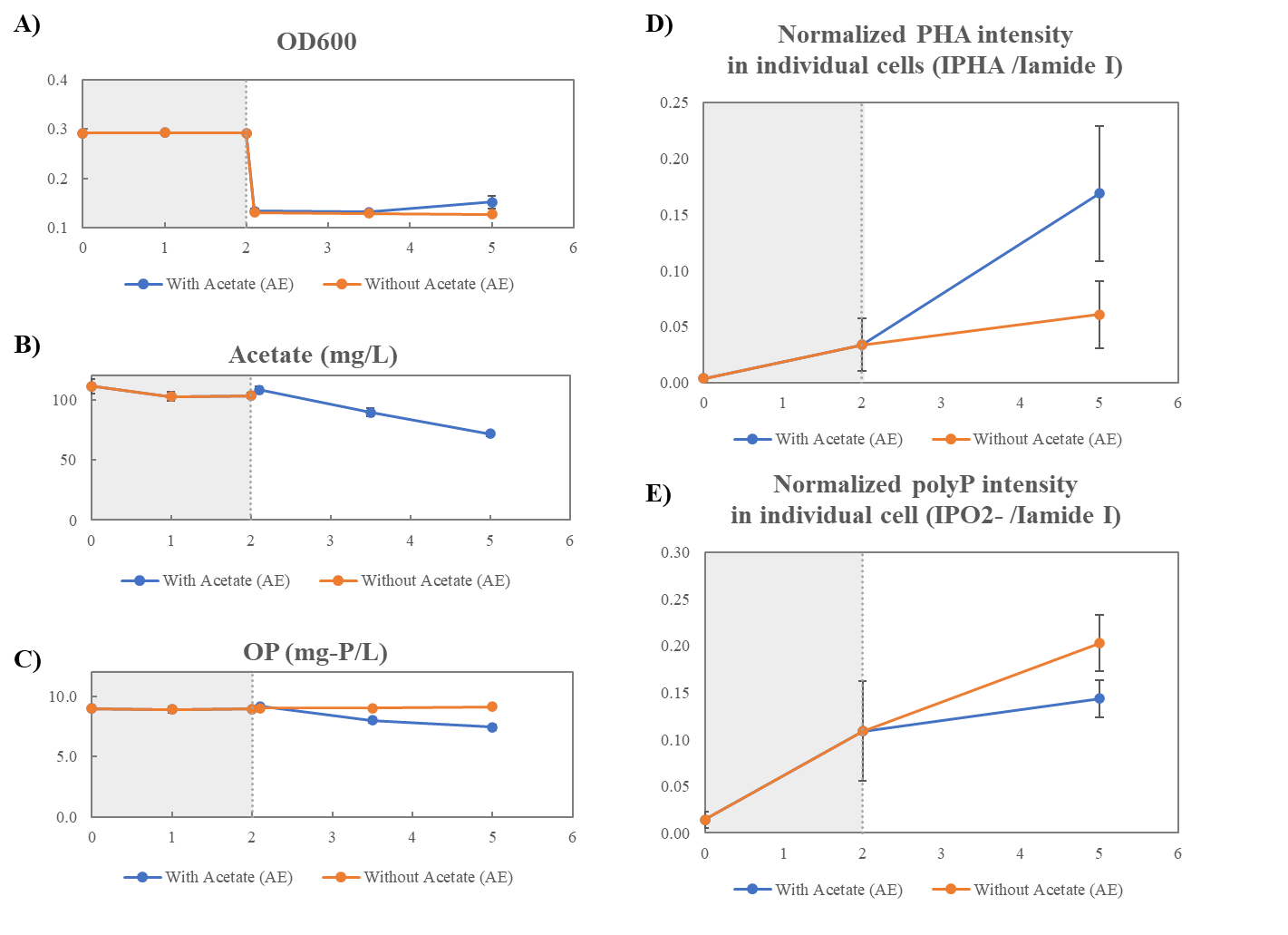

Figure S6 The variation of A) OD600, B) acetate, and C) PO_4_^3–^ concentration in the medium, and the normalized D) intracellular PHA and E) polyp intensity in individual cells in Anaerobic/Aerobic cycle analysis of pure-cultured *Acinetobacter junii*. Grey shadows indicate anaerobic phases.

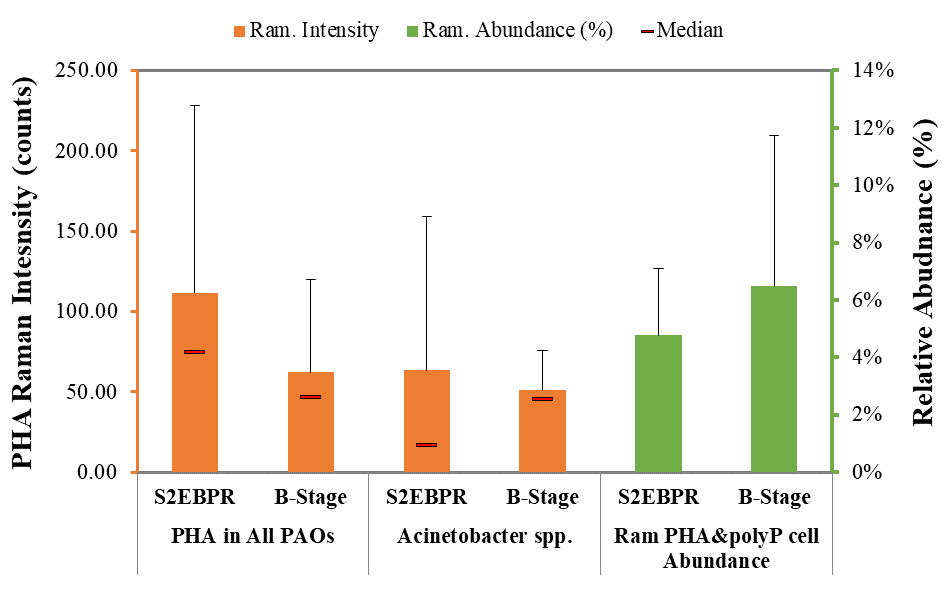

Figure S7 Average PHA signals in the polyP containing cells identified by normal SCRS and in the *Acinetobacter* identified by FISH-SCRS (orange bars), and the SCRS-based phenotypic abundance (green bars).

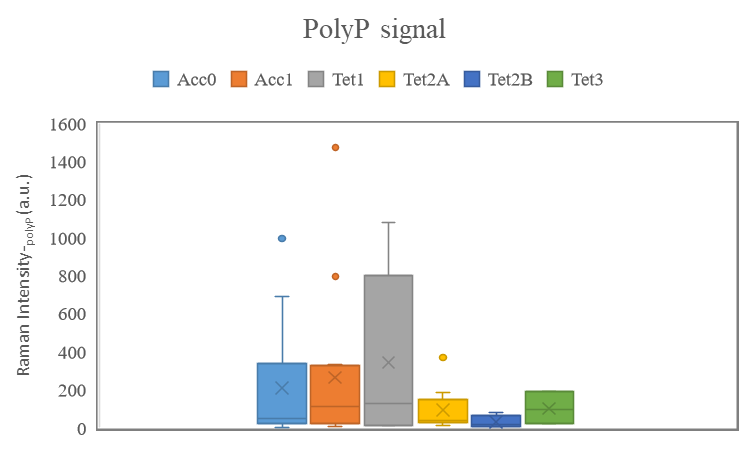

Figure S8 Average PolyP signals in *Acinetobacter*, *Accumulibacter* and *Tetrasphaera* identified by FISH-SCRS (blue bars),

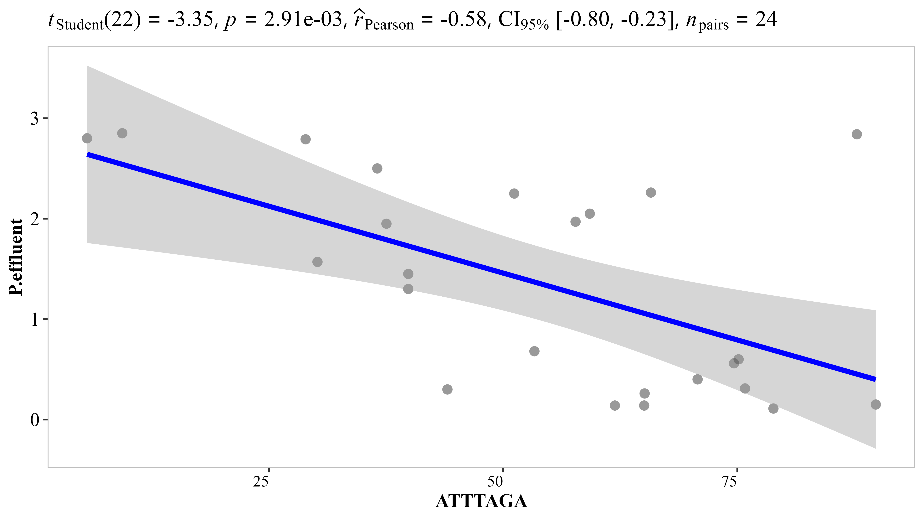

Figure S9 Linear correlation between the relative abundance of oligo1 (ATTTAGA) and the effluent-P concentrations (mg-P/L)

Table S 1. Influent and effluent PO_4_^-^-P of the P(D)N-S2EBPR system, and other full-scale EBPR and S2EBPR facilities. (Unit:)

| Process/Period  Measurement | P(D)N-S2EBPR | | | |  | S2EBPR^*^ | | | EBPR*^⁑^* |
| --- | --- | --- | --- | --- | --- | --- | --- | --- | --- |
|  | 1 – 149 | 150 – 267 | | 268 – 313 |  | SC-SSR | WR-SSRC | CC-SSM |  |
| Inf. sCOD (mg/L) | 94.0 ± 19.5 | 84.5 ± 16.1 | | 105.5 ± 21.1 |  |  |  |  | 248-425.5 |
| Eff. sCOD (mg/L) | 29.2 ± 6.7 | 23.5 ± 5.2 | | 40.4 ± 3.6 |  |  |  |  |  |
| Inf. BOD (mg/L) | 77.3 ± 29.8 | 63.6 ± 9.9 | | 89.8 ± 18.4 |  | 284.4 ± 69 | 240 ± 53 | 236 ± 92 | 148-199 |
| Inf. TP (mg-P/L) | 3.6 ±0.6 | 3.2 ±0.5 | | 4.4 ±0.6 |  | 7.1 ± 0.1 | 6.8 ± 1.0 | 2.7 ± 5.4 | 5.5-14.6 |
| Inf. PO_4_^−^-P (mg-P/L) | 2.7 ± 0.4 | 3.2 ± 0.7 | | 3.8 ± 0.4 |  |  | 3.4 | 1.8 | 3.4-10.2*^*^* |
| Eff. PO_4_^−^-P (mg-P/L) | 1.5 ± 1.0 | 0.9 ± 0.6 | | 2.4 ± 1.2 |  | 0.4 ± 0.4 | 0.2 ± 0.1 | 0.9 ± 0.3 | 0.1-0.6*^*^* |
| 3.84% | 0.1 | 0.1 | | 0.6 |  | 0.10 | 0.01 | 0.10 | - |
| 50% | 1.4 | 0.6 | | 2.3 |  | 0.28 | 0.04 | 0.82 | 0.26 |
| 90% | 2.8 | 2.1 | | 3.6 |  | 0.89 | 0.10 | 1.10 | 1.6 |
| sCOD/P (g COD/g P) | 28.2 ± 8.5 | 28.3 ± 5.9 | | 32.1 ± 10 |  |  | 23.6 |  |  |
| BOD/TP (g/g ) | 22 ± 9.3 | 18.6 ± 3.5 | | 20.2 ± 2.3 |  | 39 ± 6.6 | 38.4 ± 23 | 102 ± 88 | 12.6-31.9 |
|  | | | Cumulative-relative frequency distribution of efﬂuent PO_4_^-^-P | | | | | | |
| < 0.5 | 17% | 38% | | 3% |  | 73% | 99% | 23% | 24-95% |
| < 1.0 | 35% | 68% | | 9% |  | 91% | 100% | 79% | 64-99% |
| < 1.5 | 38% | 80% | | 18% |  | 96% | 100% | 100% | - |
| < 2.0 | 73% | 89% | | 42% |  | 99% | 100% | 100% | 85-100% |

^*^Onnis‐Hayden et al., 2019; SC-SSR: South Cary Facility, North Carolina – Sidestream RAS Fermentations; WR: Westside Reginal Facility, British Colombia – Sidestream RAS Fermentations with carbon addition; CC – Cedar Creek Facility, Kansas – Sidestream MLSS Fermentation. ^⁑^ (Neethling, 2006)

Table S 2. *ex-situ* P-uptake and P-release tests for the P(D)N-S2EBPR system and other full-scale EBPR and S2EBPR facilities.

| Process/Period  Measurement | P(D)N-S2EBPR (day) | | | | |  | S2EBPR^†^ | | | EBPR^†^ |
| --- | --- | --- | --- | --- | --- | --- | --- | --- | --- | --- |
|  | 1 - 149^*^ | 150 – 267^*^ | 268 – 313^*^ | 226 d(HAc)^⁑^ | 226 d(Fmt) |  | SC-SSR | WR-SSRC | CC-SSM |  |
| P-rel. (mg-P g_VSS_^-1^ h^-1^) | 2.9 ± 0.8 | 5.1 ± 1.5 | 4.6 ± 2.4 | 4.81 | 3.50 |  | 5.30 | 5.90 | 4.40 | 11.5 ± 6.7 |
| P-up. (mg-P g_VSS_^-1^ h^-1^) | 1.6 ± 0.7 | 2.5 ± 0.9 | 1.9 ± 1.0 | 2.57 | 1.90 |  | 2.60 | 2.40 | 1.60 | 4.2 ± 2.8 |
| HAc up. (mg-HAc g_VSS_^-1^ h^-1^) | 29.2 ± 11.4 | 29.8± 8.7 | 25.7 ± 17.0 | 26.51 | 16.0^A^ |  | 13.50 | 17.40 | 7.68 | 26.6 ± 8.3 |
| P-rel./P-up. | 2.4 ± 1.7 | 2.1 ± 0.9 | 2.4 ± 0.6 | 1.88 | 1.85 |  | 2.0 | 2.40 | 3.20 | 3.2 ± 1.2 |
| P/HAc (mol-P/mol-C) | 0.10 ± 0.03 | 0.17 ± 0.03 | 0.23 ± 0.16 | 0.18 | 0.21^A^ |  | 0.39 | 0.38 | 0.54 | 0.4 ± 0.2 |
| Gly/HAc (mol-C/mol-C) |  |  |  | 0.50 | 0.34^B^ |  | 0.15 | 0.33 | 0.35 | 0.3 ± 0.25 |
| PHA/HAc (mol-C/mol-C) |  |  |  | 0.34 | 0.25^B^ |  | 0.72 | 0.61 | 0.80 | 0.6 ± 0.6 |
| P/PHA (mol-P/mol-C) |  |  |  | 0.26 | 0.43^B^ |  | 0.52 | 0.69 | 0.73 | 0.7 ± 0.5 |
| Gly/PHA |  |  |  | 5.86 | 1.84^B^ |  | 0.90 | 0.75 | 1.07 | 0.5 ± 0.3 |

^*^ Conversion factor 1.07 gCOD/gHAc (Lettinga et al. 1999); ^⁑^ Assuming only HAc is used for PHA and Gly formation; ^A^Considering only HAc fraction of the fermentate;

^B^ Considering PHB/PHV ratio (0.33) at the end of anaerobic phase; ^†^Onnis‐Hayden et al., 2019; SC-SSR: South Cary Facility, North Carolina – Sidestream RAS Fermentations; WR: Westside Reginal Facility, British Colombia – Sidestream RAS Fermentations with carbon addition; CC – Cedar Creek Facility, Kansas – Sidestream MLSS Fermentate

Table S 3 Diversity indices of 16S rRNA gene amplicon of the P(D)N-S2EBPR system in different periods

| Period  Diversity Index | 1 – 149 d | 150 – 267 d | 268 – 313 d |
| --- | --- | --- | --- |
| Richness (R) | 10.1 ± 1.7 | 8.6 ± 1.2 | 7.4 ± 1.4 |
| Shannon Index (H') | 5.1 ± 0.8 | 4.7 ± 0.6 | 4.5 ± 0.2 |

Table S4. Regression analysis of the abundance of *Acinetobacter* and *Tetrasphaera* (%) to the anaerobic phosphorus release rate and aerobic phosphorus uptake rate (mg-P g_VSS_^-1^ h^-1^).

|  | *Acinetobacter* | | *Tetrasphaera* | |
| --- | --- | --- | --- | --- |
| Regression Statistics | Anaerobic Release | Aerobic Uptake | Anaerobic Release | Aerobic Uptake |
| Multiple R | 0.57008 | 0.338113 | 0.322899 | 0.4861 |
| R Square | 0.324991 | 0.11432 | 0.104264 | 0.236293 |
| Adjusted R Square | 0.282803 | 0.058965 | 0.04828 | 0.188562 |
| Standard Error | 1.414954 | 0.848067 | 1.629962 | 0.787508 |
| Observations | 18 | 18 | 18 | 18 |
| *p*-value | 0.013507 | 0.169961 | 0.191234 | 0.040818 |

Table S 5. Summary of poly-P Raman signal of samples from the SBPR, B-stage reactor and FISH-Raman analysis of *ex-situ* P-release and P-uptake test. (Units: Count)

| Process  poly-P signal | S2EBPR | B-Stage | PAO activity test ACA23a | | |
| --- | --- | --- | --- | --- | --- |
|  |  |  | t = 0 min | t = 60 min | t = 120 min |
| Average | 376 ± 717 | 652 ± 756 | 57 ± 80 | 28 ± 28 | 135 ± 185 |
| Maximum | 2,933 | 4,376 | 272 | 77 | 356 |
| Median | 42 | 358 | 25 | 19 | 92 |
| Minimum | 11 | 10 | 14 | 15 | 21 |

**Supplementary materials and methods**

Pure culture analysis of *Acinetobacter junni*

*Acinetobacter junni* strain (DSM 14968) was purchased from the DSMZ Cell Lines Bank (Braunschweig, Germany) It was selected based on the similarity to our *Acinetobacter*-like OTU sequence. In these tests, fresh *A. junni* was harvested xx and transferred into 50 ml anaerobic serum bottle. Anaerobic periods last for 2 hours with 100 mg/L acetate and 10 mg-P/L PO_4_^3−^. After that, cells were washed to remove external carbon cells and split into two different media. One bottle has acetate the same as anaerobic medium, and another has no acetate in medium. The two bottles were aerated for another 3 hours. Supernatants were taken every hour to measure acetate and PO_4_^3−^ concentration. Cell samples were taken at time 0, 2, and 5 hours to measure intracellular polyhydroxyalkanoate (PHA) and poly-phophorus (polyP) via Raman spectroscopy.

**References**

Neethling, J.B., 2006. Factors Influencing the Reliability of Enhanced Biological Phosphorus Removal. IWA Publishing.

Onnis-Hayden, A., Srinivasan, V., Tooker, N.B., Li, G., Wang, D., Barnard, J.L., Bott, C., Dombrowski, P., Schauer, P., Menniti, A., Shaw, A., Stinson, B., Stevens, G., Dunlap, P., Takács, I., McQuarrie, J., Phillips, H., Lambrecht, A., Analla, H., Russell, A., Gu, A.Z., 2020. Survey of full-scale sidestream enhanced biological phosphorus removal (S2EBPR) systems and comparison with conventional EBPRs in North America: Process stability, kinetics, and microbial populations. Water Environment Research 92, 403–417.

Stokholm-Bjerregaard, M., McIlroy, S.J., Nierychlo, M., Karst, S.M., Albertsen, M., Nielsen, P.H., 2017. A Critical Assessment of the Microorganisms Proposed to be Important to Enhanced Biological Phosphorus Removal in Full-Scale Wastewater Treatment Systems. Frontiers in Microbiology 8.
